# Identifying SARS-CoV-2 Antiviral Compounds by Screening for Small Molecule Inhibitors of Nsp13 Helicase

**DOI:** 10.1101/2021.04.07.438808

**Authors:** Jingkun Zeng, Florian Weissmann, Agustina P. Bertolin, Viktor Posse, Berta Canal, Rachel Ulferts, Mary Wu, Ruth Harvey, Saira Hussain, Jennifer C. Milligan, Chloe Roustan, Annabel Borg, Laura McCoy, Lucy S. Drury, Svend Kjaer, John McCauley, Michael Howell, Rupert Beale, John F.X Diffley

**Author notes:** contributed equally (Jingkun Zeng, Florian Weissmann, Agustina P. Bertolin).

## Abstract

The coronavirus disease 2019 (COVID-19) pandemic, which is caused by severe acute respiratory syndrome coronavirus 2 (SARS-CoV-2) is a global public health challenge. While the efficacy of vaccines against emerging and future virus variants remains unclear, there is a need for therapeutics. Repurposing existing drugs represents a promising and potentially rapid opportunity to find novel antivirals against SARS-CoV-2. The virus encodes at least nine enzymatic activities that are potential drug targets. Here we have expressed, purified and developed enzymatic assays for SARS-CoV-2 nsp13 helicase, a viral replication protein that is essential for the coronavirus life cycle. We screened a custom chemical library of over 5000 previously characterised pharmaceuticals for nsp13 inhibitors using a FRET-based high-throughput screening (HTS) approach. From this, we have identified FPA-124 and several suramin-related compounds as novel inhibitors of nsp13 helicase activity *in vitro*. We describe the efficacy of these drugs using assays we developed to monitor SARS-CoV-2 growth in Vero E6 cells.

## Introduction

More than 100 million people have been infected and more than 2.5 million people have died worldwide due to the COVID-19 pandemic as of late February 2021 (1). The number of infections continues to rise, with more than 150,000 daily infections globally, highlighting the importance of developing effective vaccines and therapeutics for the prevention and treatment of COVID-19. Vaccines are under mass roll-out nevertheless concerns over potentially vaccine-resistant variants have been growing. It has been shown in cell culture experiments that the virus has the potential to evolve mutant strains that can evade neutralizing antibodies produced by the body to combat infections (2), and reports have emerged that certain variants could cause re-infections (3). Many of the vaccines being used, e. g. the COVID-19 vaccines produced by Pfizer, Moderna and AstraZeneca-Oxford, target the spike protein on the surface of SARS-CoV-2 (4). Multiple mutations of the spike protein have been found around the globe (5). Although there is no clear evidence of vaccine-evading variants yet, vaccines against rapidly evolving structural proteins might not protect against all newly emerging strains and are unlikely to have pan-coronavirus efficacy. With these uncertainties regarding SARS-CoV-2 vaccines, multiple layers of protection and treatment against COVID-19 are needed including the identification of therapeutic drugs that can interfere with viral entry or viral propagation is of utmost importance. Nonetheless, therapeutic options for COVID-19 are currently limited. *De novo* development of antiviral therapies generally requires between 10 to 17 years (6). The repurposing of drugs originally developed for other uses could provide a practical approach for the fast identification, characterisation and deployment of antiviral treatments (6, 7). The repurposing of approved or investigational drugs exploits existing detailed information on drug chemistry together with human pharmacology and toxicology, allowing rapid clinical trials and regulatory review (6). This strategy has proven useful so far, with a few repurposed medicines having been authorised by different regulatory agencies to treat COVID-19, such as remdesivir, an antiviral developed to treat Ebola (8, 9). Remdesivir is a pro-drug inhibitor of the RNA-dependent RNA polymerase (RdRp) that shows inhibitory activity against all three strains, SARS-CoV-1, MERS-CoV and SARS-CoV-2, of the coronavirus outbreaks in this century (10–12). Drug resistance to monotherapies may develop rapidly, particularly in RNA viruses where mutations occur frequently, thus it would be useful to have multiple antiviral drugs (13, 14).

SARS-CoV-2 is a positive-sense single stranded RNA virus that encodes at least nine enzymatic activities in two overlapping large polyproteins pp1a and pp1ab (15, 16). Once expressed in the host cell, pp1a and pp1ab are processed by virus-encoded proteases into 16 non-structural proteins (nsps) (15, 16). Coronavirus nsp13, one of the non-structural proteins, is a superfamily 1B (SF1B) helicase that can unwind DNA or RNA in an NTP-dependent manner with a 5’ to 3’ polarity (17–20). Moreover, nsp13 harbours RNA 5’-triphosphatase activity that could play a role in viral 5’ RNA capping (18, 20, 21). Nsp13 is highly conserved among SARS-like coronaviruses with 99.8% sequence identity (600 out of 601 amino acids) between SARS-CoV-1 and SARS-CoV-2 (22). Importantly, nsp13 is a key component of replication-transcription complexes (RTC) and is indispensable for the coronavirus life cycle, making it a promising target for pan-coronavirus antivirals (23–26).

As part of a larger project to identify small molecule inhibitors of all SARS-CoV-2 enzymes, we report the development of a high throughput fluorescence resonance energy transfer (FRET)-based assay for nsp13 helicase activity *in vitro.* We used this assay to screen a custom compound library of over 5000 previously characterised pharmaceuticals. We identify FPA-124 and suramin-like compounds as novel nsp13 inhibitors that also show antiviral activity in a cell-based viral proliferation assay.

## Results

### Protein expression and purification

For SARS-CoV-1 nsp13, it has previously been shown that GST-tagged nsp13 expressed in insect cells is more active than MBP- or 6xHis-tagged nsp13 expressed in bacteria (27, 28). Therefore, we used a baculovirus-insect cell expression system to express and purify two differently tagged SARS-CoV-2 nsp13 versions, GST-nsp13 and 3xFlag-His_6_-nsp13 (FH-nsp13). Both nsp13 variants were purified using affinity chromatography based on GST or the 3xFlag tag followed by gel filtration (**Figure 1A** and **Supplementary Figure S1A-B**).

**Figure 1.**
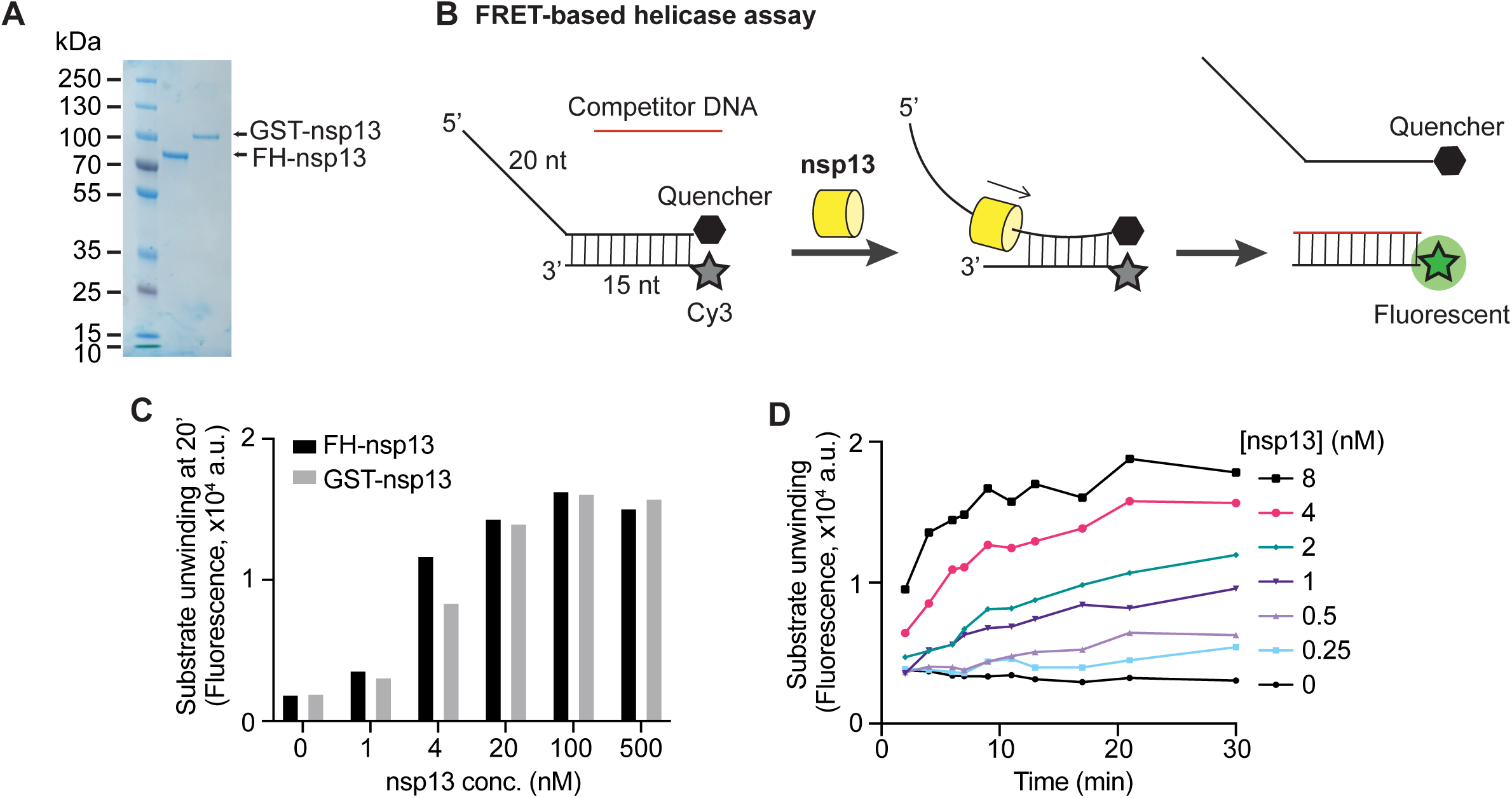
Development of a FRET-based SARS-CoV-2 nsp13 helicase assay. (**A**) Purified 3xFlag-His_6_-nsp13 (FH-nsp13) and GST-nsp13 analysed by SDS-PAGE and Coomassie staining. (**B**) Schematic diagram illustrating the FRET-based nsp13 helicase assay. A nucleic acid substrate with a 5’ overhang is generated by annealing a fluorophore-labelled oligonucleotide (Cy3 strand) with a quencher-containing oligonucleotide (BHQ-2 strand). Nsp13 helicase activity displaces the BHQ-2 strand generating a fluorescent signal. A competitor oligonucleotide captures the free Cy3 strand preventing substrate re-annealing. The generated Cy3 - competitor hybrid does not contain a 5’ overhang and is no longer a good substrate for nsp13. (**C**) FH-nsp13 and GST-nsp13 were incubated with 50 nM DNA substrate and 2 mM ATP at the indicated concentrations for 20 min on ice and Cy3 signal was measured (a.u.= arbitrary units). (**D**) Time course experiment using FH-nsp13 as indicated, 50 nM DNA substrate, and 2 mM ATP (a.u.= arbitrary units).

### Development of a FRET-based helicase assay

Coronavirus helicases unwind either double stranded DNA or RNA, translocate in a 5′-to-3′ direction and require a 5′-overhang to load on their substrate and start duplex unwinding (18-20, 27, 29). To monitor the unwinding activity of nsp13, we designed a FRET-based helicase assay suitable for high-throughput screening. The substrate consisted of a 35 nucleotide (nt) DNA strand hybridized to a second 15 nt complementary DNA, leaving a 20 nt single-stranded overhang at the 5’ end. A Cy3 fluorescent moiety and a non-emitting dark quencher (BHQ-2) were present at the 5’ and 3’ end of the duplex region, respectively, so the Cy3 would be quenched until nsp13 unwinds the strands, freeing the fluorophore from quenching and producing fluorescence (**Figure 1B**). Indeed, Cy3 fluorescence was quenched by the BHQ-2 present on the opposite strand (**Supplementary Figure S1C**), and this quenching was lost following incubation with nsp13 (**Figure 1C**). A 5-fold excess of competitor DNA strand was added to prevent substrate re-annealing (**Supplementary Figure S1D)**. This competitor strand captures one of the unwound strands (Cy3 strand) generating a DNA duplex without 5’ overhang, and hence, a very poor substrate for nsp13 (**Figure 1B**). Both enzyme preparations, FH-nsp13 and GST-nsp13, showed similar levels of DNA substrate unwinding after a 20-minute incubation on ice (**Figure 1C**). Due to the higher protein yields obtained after FH-nsp13 expression and purification compared to the GST version, we chose FH-nsp13 for all subsequent experiments. Unwinding increased approximately linearly over the first 5-10 min. of the reaction at FH-nsp13 concentrations of 1 nM-4 nM (**Figure 1D**). We also found nsp13 was not able to unwind a similar DNA substrate with a 3’ overhang (**Supplementary Figure S1E-F**), consistent with other coronavirus helicases (18–20, 30). FH-nsp13 unwinding activity was also observed in a gel-based assay using a radiolabelled DNA substrate with the same DNA sequence as the substrate used in the FRET-based assay (**Supplementary Figure S1G**).

In order to determine the optimal concentration of substrates for best sensitivity in nsp13 inhibitor screening, we determined the Michaelis-Menten constants (K_m_) for ATP and nucleic acid substrates. It has been shown by others that coronavirus helicases lack nucleic acid specificity, being capable of unwinding DNA and RNA (18–20). We synthesised an RNA substrate of identical sequence as its DNA counterpart and tested nsp13 activity. Nsp13 exhibited similar activity on both the DNA and RNA substrates, with a higher K_m_ value for the DNA (K_m_=2.6 µM, **Figure 2A** **and Supplementary Figure S2A**) than the RNA substrate (K_m_= 1.0 µM, **Figure 2B** **and Supplementary Figure S2B**). Unwinding of the RNA substrate by FH-nsp13 was also observed in a gel-based assay (**Supplementary Figure S2C**). The K_m_ values obtained for ATP were similar with either the DNA (K_m_= 0.11 mM, **Figure 2C** **and Supplementary Figure S2D**) or the RNA substrate (K_m_= 0.13 mM, **Figure 2D** and **Supplementary Figure S2F**) in the reaction. As expected, in the absence of ATP, both substrates (DNA/RNA) presented only background levels of fluorescence (**Supplementary Figure S2D,E**). The DNA substrate was used in the subsequent high throughput screen (HTS) because it is more stable and was easier to source during the UK’s first national lockdown. For HTS reactions, we chose to use an ATP concentration close to the K_m_ value of 100 µM, and a DNA substrate concentration of 180 nM capable of producing strong fluorescent signals. We used 3 nM nsp13 in HTS reactions after performing an enzyme titration around final DNA substrate and ATP concentrations (**Figure 2E**). We confirmed that nsp13 activity was not affected by DMSO at concentrations up to 5% as the chemical compounds to be screened were dissolved in DMSO (**Supplementary Figure S2F).**

**Figure 2.**
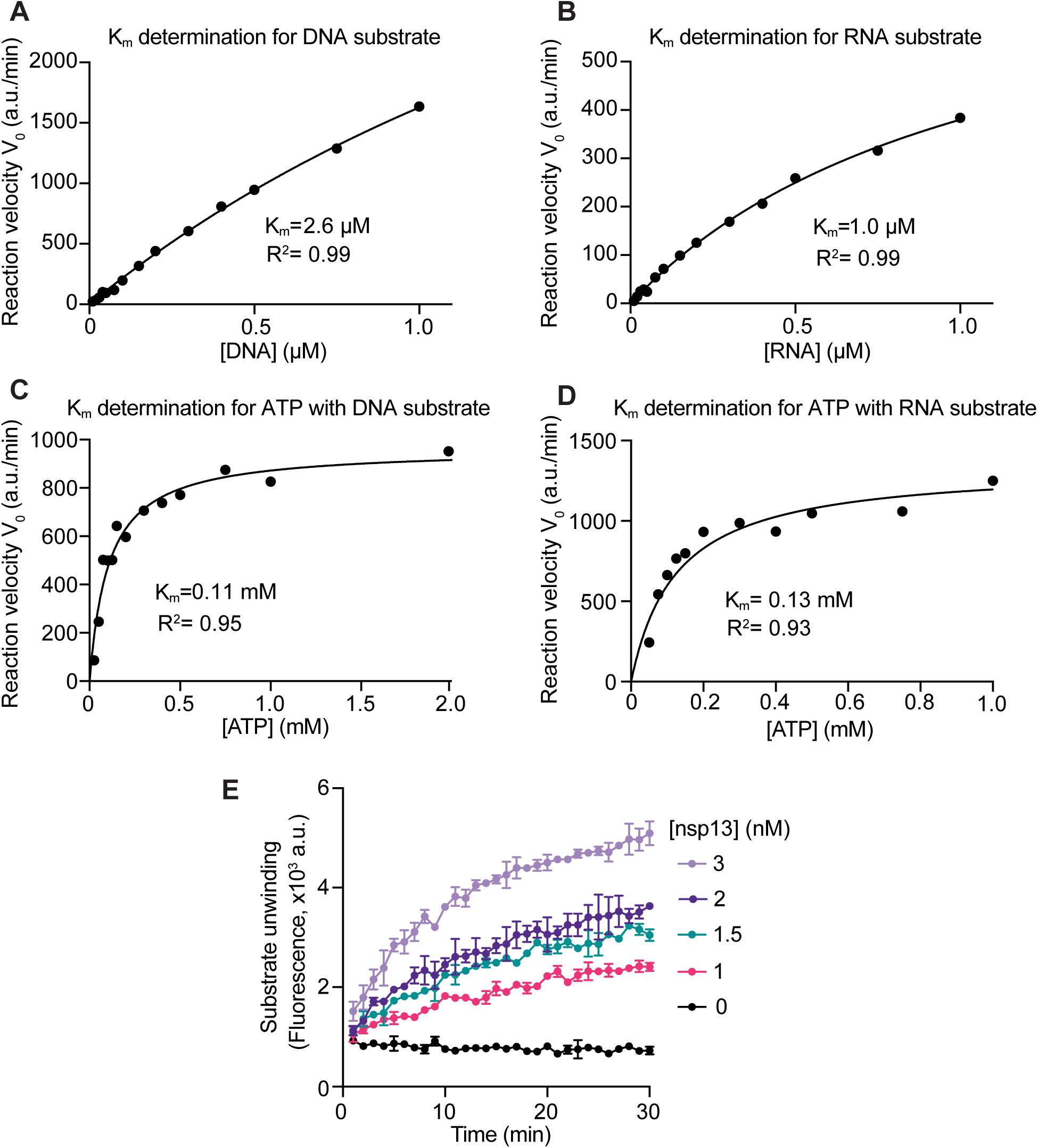
Enzyme kinetics and optimisation of assay conditions for HTS. (**A**) Michaelis-Menten kinetics (K_m_) for the DNA substrate. To obtain the Michaelis constant (K_m_) of nsp13 for DNA, initial reaction velocities (V_0_) were determined at various DNA substrate concentrations. The experiment was performed using 1 nM FH-nsp13 and 1 mM ATP (see **Supplementary Figure S2A**). The data was fitted using the Michaelis-Menten equation (a.u./min: arbitrary units per min) using Prism software. (**B**) K_m_ for the RNA substrate. The experiment was performed as in (A) using 1 nM FH-nsp13 and 1 mM ATP (see **Supplementary Figure S2B**). (**C**) K_m_ for ATP with the DNA substrate in reaction. The experiment was performed as in (A) using 2 nM FH-nsp13 and 200 nM DNA substrate (see **Supplementary Figure S2D**). (**D**) K_m_ for ATP with the RNA substrate in reaction. The experiment was performed as in (A) using 3 nM FH-nsp13 and 200 nM RNA substrate (see **Supplementary Figure S2E**). (**E**) Helicase assay at substrate concentrations suitable for HTS. A time course of Cy3 signal was recorded using the indicated FH-nsp13 concentrations, 100 µM ATP and 200 nM DNA substrate.

### Chemical library screen design and results

We performed a HTS to identify inhibitors of SARS-CoV-2 nsp13 helicase activity using a custom chemical library containing over 5000 compounds. The compounds were dispensed into 384-well plates between columns 3 and 22. Wells in columns 2 and 23 on each plate served as solvent controls containing only DMSO (**Supplementary Figure S3A**). Nsp13 was incubated with compounds for 10 min, then the substrates were dispensed to start the reaction (**Figure 3A**). The screen was performed at two compound concentrations, 1.25 and 6.25 µM. We observed that, on the day of the screen, the reaction was complete by 10 minutes (**Supplementary Figure S3B**). The time required to dispense substrates to all the wells within a plate to start the reaction was ∼10 seconds, while the time required to read all wells was ∼75 seconds (**Supplementary Figure S3C-D**). We applied a positional correction to the calculation of the initial velocity of reactions for each plate to compensate for this time delay (**Figure 3B-C**). A total of 339 compounds showed >20% reduction of corrected initial velocity or showed >10% reduction of endpoint signals at either 1.25 µM or 6.25 µM compound concentration. After manual inspection of their kinetic curves, 142 of the 339 compounds showed a clear effect in HTS reactions and were selected as primary hits (example in **Figure 3D**). To find specific nsp13 inhibitors, we initially selected 35 of the primary hits that displayed >30% reduction of corrected initial velocity for subsequent validation experiments. (**Figure 3E**).

**Figure 3.**
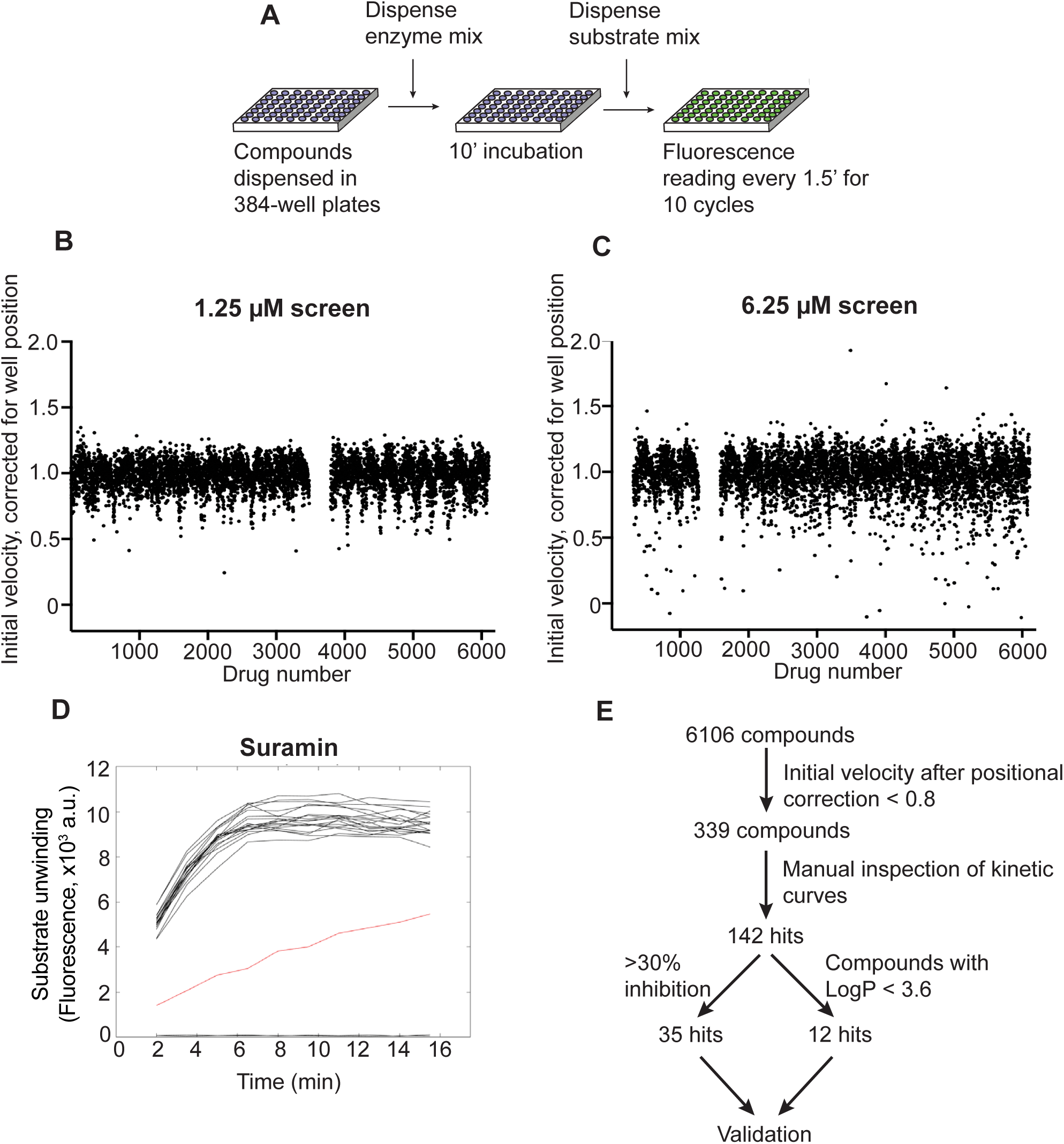
High-throughput screening for inhibitors of SARS-CoV-2 nsp13 helicase. (**A**) Logistics of the HTS. A custom compound library was screened in 384-well plate format for inhibitors of nsp13 using the FRET-based helicase assay. A nsp13 solution was dispensed into compound-containing plates. After 10 minutes, reactions were started by addition of a substrate solution and fluorescence readings were taken in 90 s intervals. (**B, C**) Results of the screens performed at 1.25 µM (B) and 6.25 µM (C) compound concentration. Initial reaction velocity was corrected for well-position and plotted against compound number. (**D**) Kinetic curves of compounds that showed at least 20% reduction in corrected velocity were inspected manually. As example, time-course data for the compound suramin in the 6.25 µM screen is shown (red curve, data from 20 surrounding wells in black). (**E**) Summary of HTS results. The process of hit selection for further validation is shown.

### Characterisation of HTS hits

In a first validation experiment, we tested the compounds using the same assay conditions as in the HTS and determined the compound concentration at which half maximal nsp13 inhibition (IC_50_) was observed. We found that 27 of the 33 tested compounds showed nsp13 inhibition with apparent IC_50_ values < 30 µM (**Supplementary Figure S4** and **Table 1**, second column). We then characterised the hits under more physiological conditions where the DNA substrate was replaced by its RNA counterpart, and the ATP concentration was increased from 100 µM to 2 mM. Nonspecific inhibition of enzymes due to colloidal aggregation of compounds is the most common source of false-positives in high-throughput inhibitor screens (31–34). The presence of non-ionic detergents can reduce colloid formation, which can lead to a right-shift of dose-response curves for many aggregation-prone compounds (35). In order to uncover potential aggregators, we added 0.02% Tween-20 to the reaction. Surprisingly, under these conditions only 5 out of 27 compounds still inhibited nsp13 and showed similar IC_50_ values as in the first validation experiment (**Supplementary Figure S5** and **Table 1**, third column). The remaining 22 compounds lost the strong inhibition shown in the first experiment, as indicated by the increase in their IC_50_ values (**Table 1**, compare second column versus third column).

**Table 1.**
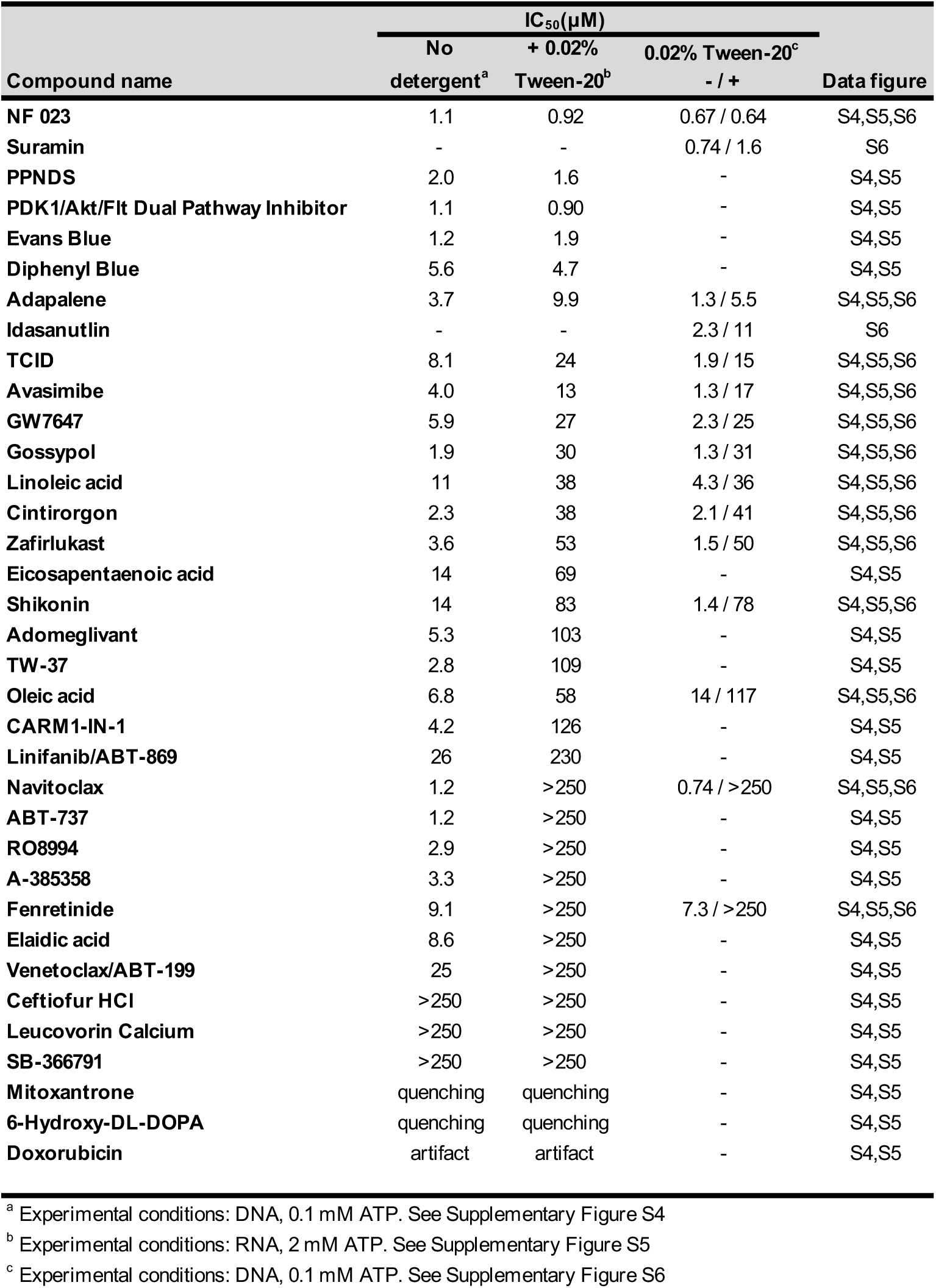
Inhibitory activity and detergent-sensitivity of 35 identified compounds against SARS-CoV-2 nsp13

To test directly if some of our compounds could inhibit nsp13 activity via a detergent-sensitive aggregation mechanism (32, 35), we tested 15 selected compounds in the presence or absence of Tween-20 with otherwise identical assay conditions (**Supplementary Figure S6**). The presence of Tween-20 reduced the inhibition of 13 of the 15 tested compounds, causing an IC_50_ shift greater than 4-fold (**Supplementary Figure S6** and **Table 1**, fourth column), suggesting the nsp13 inhibition shown by these drugs in HTS and in the first validation experiment could be due to an aggregation effect. Colloidal aggregators absorb and inhibit enzymes without specificity (31, 33). Potential aggregators can be further confirmed by counter-screening against other enzymes (33). Hence, we tested whether these compounds could inhibit the SARS-CoV-2 RNA-dependent RNA polymerase (RdRp) using a FRET-based assay developed in an accompanying manuscript (Bertolin et al.). Compounds with mild detergent sensitivity that still showed moderate to weak nsp13 inhibition in the presence of detergent, also inhibited RdRp in a similar fashion (e.g., Avasimibe, **Supplementary Figure S7** and **Supplementary Table S1**). On the other hand, compounds that no longer inhibited nsp13 in the presence of detergents did not inhibit RdRp (e.g., A-385358, **Supplementary Figure S7** and **Supplementary Table S1**). All together, these results suggest that many of the compounds identified in the nsp13 HTS were aggregation-based inhibitors. These findings are consistent with Aggregator Advisor, an open access tool that advises on the likelihood of colloidal aggregation based on criteria including lipophilicity (LogP>3.5) and on a structural similarity index (>85% similarity compared to reported aggregators) (36). We analysed the selected hits using this method and, not surprisingly, 21 out of 22 detergent-sensitive compounds showed LogP>3.5 (**Supplementary Table S2**).

After this first round of analysis, we re-examined our primary hit list, and selected another 12 hit compounds with a LogP<3.6 and with no significant similarity to known aggregators (**Supplementary Table S3** and **Figure 3E**) (36). We also tested two published nsp13 inhibitors, SSYA10-001 (37, 38) and myricetin (39). We determined the IC_50_ values of these compounds in the presence or absence of Tween-20 (**Table 2** and **Supplementary Figure S8**). Eight of them did not show clear nsp13 inhibition in any condition. Hypocrellin A, TAK875 and elvitegravir together with the published nsp13 inhibitor myricetin showed clear inhibition only in the absence of detergent (**Table 2** and **Supplementary Figure S8**). The other published nsp13 inhibitor SSYA10-001 presented moderate sensitivity to detergent. FPA-124 was the only compound that inhibited nsp13 with no detergent sensitivity in this test (**Table 2** and **Supplementary Figure S8-S9**).

**Table 2.** Inhibitory activity against SARS-CoV-2 nsp13 and detergent-sensitivity of 12 HTS hits selected in the second validation round and 2 control compounds

| Compound name | IC<sub>50</sub> (μM) |  |  | Data figure |
| --- | --- | --- | --- | --- |
|  | No detergent <sup>a</sup> | + 0.02% | + 0.01% Triton |  |
|  |  | Tween-20 <sup>a</sup> | X-100 <sup>b</sup> |  |
| FPA 124 | 8.5 | 8.4 | 8.9 | S8,S9 |
| Hypocrellin A | 12 | >250 | - | S8 |
| TAK875 | 19 | 72 | - | S8 |
| Elvitegravir | 22 | >250 | - | S8 |
| NDP-BVU-972 | 75 | >250 | - | S8 |
| Eplerenone | >250 | no inhibition | - | S8 |
| Fluocinolone Acetonide | >250 | no inhibition | - | S8 |
| Omapatrilat | >250 | no inhibition | - | S8 |
| ADL5859 | >250 | no inhibition | - | S8 |
| Nedocromil | no inhibition | no inhibition | - | S8 |
| Lifitegrast | no inhibition | no inhibition | - | S8 |
| SKF 89145 | no inhibition | no inhibition | - | S8 |
| Suramin <sup>c</sup> | 0.94 | 1.1 | 1.6 | S8,S9 |
| Zafirlukast <sup>c</sup> | 6.6 | 55 | 22 | S8,S9 |
| SSYA10-001 <sup>d</sup> | 7.5 | 28 | 21 | S8,S9 |
| Myricetin <sup>e</sup> | 24 | 125 | - | S8 |

<sup>a</sup> Experimental conditions: RNA, 0.1 mM ATP. See Supplementary Figure S8
<sup>b</sup> Experimental conditions: RNA, 0.1 mM ATP. See Supplementary Figure S9
<sup>c</sup> Suramin and zafirlukast, compounds characterised in Table 1, were included for comparison purposes.
<sup>d</sup> Literature compound. See (Adedeji et al., Antimicrob Agents Chemother, 2012)
<sup>e</sup> Literature compound. See (Yu et al., Bioorg Med Chem Lett, 2012)

After all the above analyses, we found 7 compounds that inhibited nsp13 with no detergent-sensitivity (PDK1/Akt/Flt Dual Pathway Inhibitor, suramin and suramin-related compounds, and FPA-124, **Supplementary Figure S10**). PDK1/Akt/Flt Dual Pathway Inhibitor showed promiscuous inhibition in several other screens, including the RdRp screen, nsp3 papain-like protease screen, nsp5 main protease screen and nsp15 endonuclease screen (Biochem J, this issue). Therefore, it was excluded from further analysis. Suramin and suramin-related compounds (NF 023, PPNDS, Evans Blue and Diphenyl Blue) are heavily negatively charged molecules and may inhibit nucleic acid-binding proteins by binding to positively charged protein regions (40–44). They were also identified as hits in the RNA-dependent RNA polymerase (RdRp) screen (Bertolin et al. Biochem J, this issue) but not in other parallel screens, consistent with the ability of these polyanionic compounds to strongly inhibit certain nucleic acid-binding proteins (41, 45–47). Notably, a kinase inhibitor, FPA-124, inhibited nsp13 with a micromolar IC_50_ of ∼9 µM, and its IC_50_ value was not affected by the addition of the non-ionic detergents Tween-20 or Triton X-100 (**Supplementary Figures S8-S9**), suggesting its nsp13 inhibition was unlikely to be due to aggregation. Moreover, FPA-124 did not score as a hit in the RdRp screen nor parallel screens against other SARS-CoV-2 enzymes (Biochem J, this issue), indicating its inhibitory activity is likely target specific. Examples of the observed IC_50_ curves are shown in **Figure 4**. Suramin, NF 023 and FPA-124 are compounds whose enzymatic inhibition is largely insensitive to detergent addition (**Figure 4A-C**). On the other hand, navitoclax and linoleic acid are examples of compounds with high LogP values and strong sensitivity to detergents suggesting that their inhibitory effect could be attributed to colloidal aggregation occurring at certain micromolar concentrations (**Figure 4D-E**).

**Figure 4.**
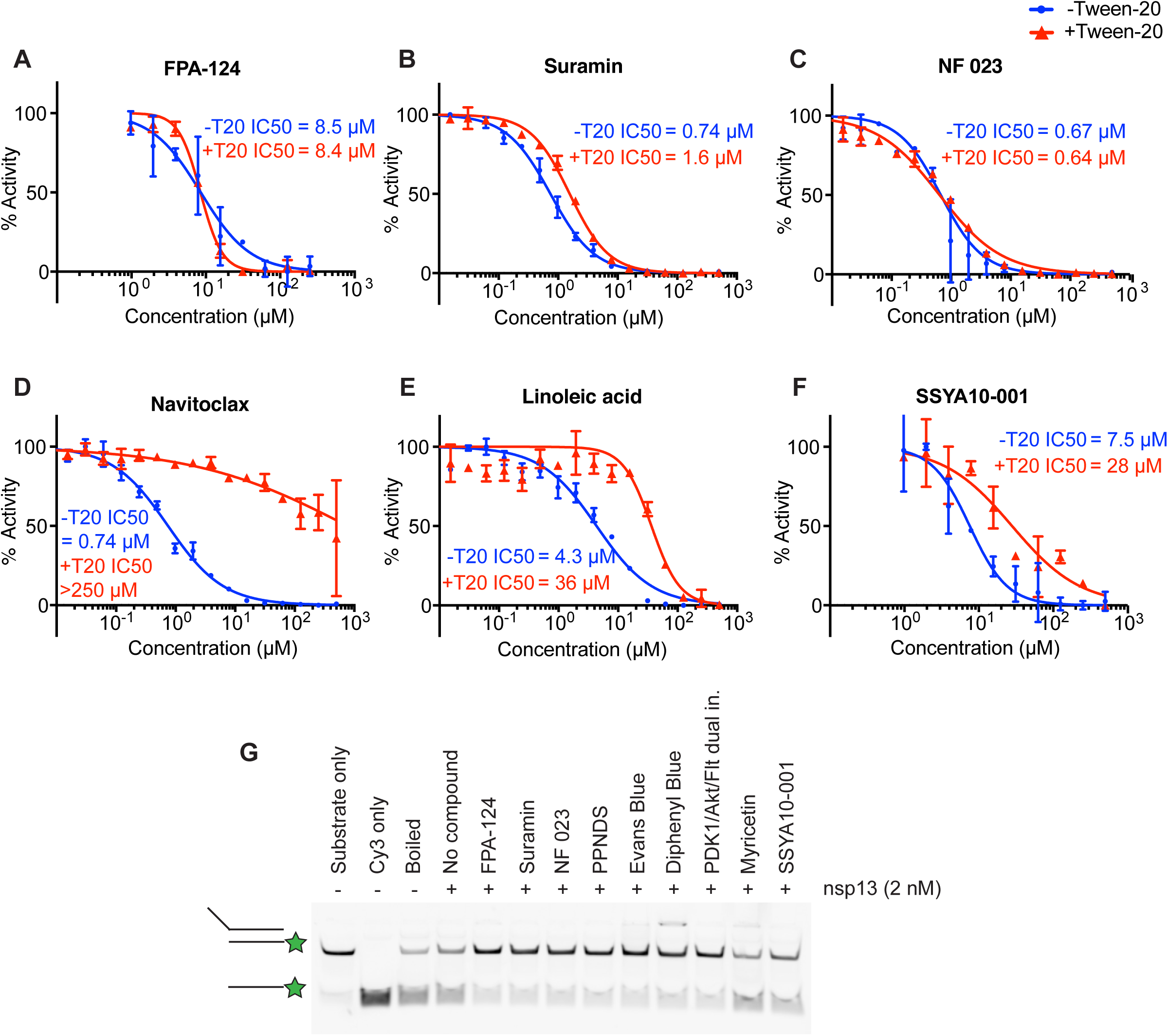
*In vitro* validation of selected compounds identified as nsp13 inhibitors. **(A-F)** Concentration-response curves of selected compounds determined using the FRET-based nsp13 helicase assay. The results shown here are part of the larger experiment shown in **Supplementary Figure S6** (for B, C, D, E) or **S7** (for A, F). The experiment was performed in the presence (+T20) or absence (-T20) of 0.02% Tween-20. FPA-124 (**A**), Suramin (**B**) and NF 023 (**C**) showed little or no detergent sensitivity, whereas Navitoclax (**D**), Linoleic acid (**E**) and SSYA10-001 (**F**) showed an increase in IC_50_ when Tween-20 was included. IC_50_ values were calculated with Prism software. (**G**) Validation of selected compounds in a gel-based nsp13 helicase assay visualising inhibition of duplex unwinding. Compounds were incubated with 2 nM FH-nsp13 at 15 µM compound concentration for 10 min before reactions were started by addition of 2 mM ATP and 50 nM RNA substrate. After 5 min reaction products were analysed by native PAGE and visualisation of Cy3 fluorescence. Controls: RNA substrate (Substrate only), Cy3 strand only (Cy3 only), RNA substrate after 5 min incubation at 95 °C (Boiled).

We also tested these compounds in a gel-based nsp13 helicase assay (**Figure 4G**). We used an RNA substrate consisting of a Cy3 strand and an unlabelled complementary strand with a 5’ overhang. A competitor strand was included to capture the unlabelled strand. Reaction products were then separated by native PAGE and detected by Cy3 fluorescence. We tested the identified detergent-insensitive compounds -FPA-124, suramin, NF 023, PPNDS, Evans Blue, Diphenyl Blue together with the 2 published SARS-CoV-1 nsp13 inhibitors, myricetin and SSYA10-001, at a compound concentration of 15 µM in the presence of 0.01% Triton X-100. All 6 identified compounds clearly reduced the generation of single-stranded Cy3 strand, confirming that the compounds inhibited helicase activity (**Figure 4G**). As expected, based on their IC_50_ values in the presence of detergent (**Table 2**), myricetin and SSYA10-001 inhibited nsp13 less efficiently under these conditions.

### Viral inhibition assays

We next evaluated potential antiviral activity of the compounds against SARS-CoV-2 in Vero E6 cells. Our experiments thus far identified 6 compounds that inhibited nsp13 in biochemical assays. Of these, suramin, PPNDS, NF 023, Evans Blue and Diphenyl Blue are structurally related compounds containing at least one polysulfonated naphthyl group, which is believed to be the critical pharmacophore (40, 41). Suramin is the prototypical naphthalene polysulfonated compound and most studied drug of this group, so we tested suramin in our cell-based experiments. The other validated inhibitor was FPA-124, which is a cell permeable selective AKT inhibitor (48). We also included the two published SARS-CoV-1 nsp13 inhibitors SSYA10-001 and myricetin (37–39). To test an effect on viral replication, the compounds were added to Vero E6 cells, and then cells were infected with SARS-CoV-2 at a multiplicity of infection of 0.5. After 22 hours cells were fixed and analysed by immunofluorescence (**Figure 5A**). Immunofluorescent detection of SARS-CoV-2 N protein was used as a read out for viral replication in cell culture (see Material and Methods). The compounds were tested over a range of concentrations, and the half maximal effective concentration (EC_50_) for each compound was calculated (**Figure 5B-C**). Suramin and FPA-124 showed viral inhibition with EC_50_ values of 9.9 µM and 14 µM, respectively, while the published nsp13 inhibitors, myricetin and SSYA10-001, presented a weaker viral inhibition potency (EC_50_=32 µM and 81 µM, respectively). Of these four tested compounds, only FPA-124 showed considerable inhibition of cellular growth at high concentrations (100–300 µM), perhaps due to inhibition of AKT kinase (**Supplementary Figure S11A**) (48).

**Figure 5.**
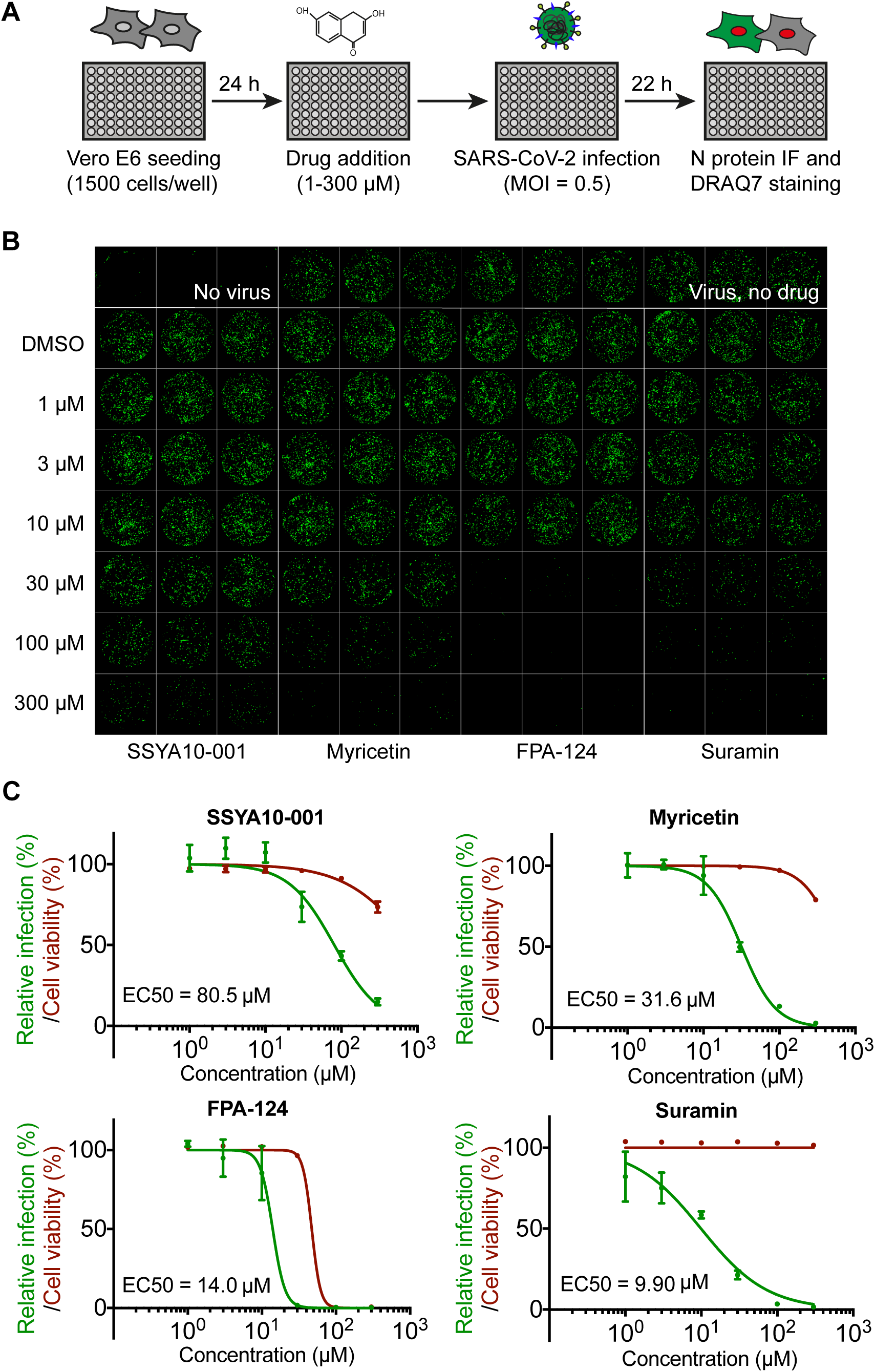
Antiviral activity of selected compounds against SARS-CoV-2 in cell culture. (**A**) Schematics of the viral infection inhibition protocol. In brief, Vero E6 cells were seeded in a 96-well format and after 24 hours were treated with selected compounds at defined concentrations. The cultures were infected with SARS-CoV-2 at a MOI of 0.5 PFU/cell. Twenty-two hours after infection, cells were fixed and analysed by immunofluorescence staining and imaging. (**B**) Confocal microscope images of viral N protein staining (green) show anti-SARS-CoV-2 activities of SSYA10-001, myricetin, suramin and FPA-124 at indicated concentrations. The corresponding DRAQ7-stained cell nuclei images are shown in **Supplementary Figure S11A**. (**C**) Dose-response curve analysis by immunofluorescence for each drug. Viral infection was measured as the area of viral plaques stained for N protein (green lines) and cell viability as the area of cells stained for DRAQ7 (red lines). Data is plotted as percentage compared to DMSO only control wells (100%). Values represent mean and standard deviation (SD) of 3 replicates. Areas were determined using FIJI software and EC_50_ values were calculated with Prism software.

Combining two drugs with antiviral activity and different viral or cellular targets can result in improved outcomes compared to antiviral monotherapy (49). Therefore, we decided to investigate potential synergy between nsp13 inhibitors and the RNA-dependent RNA polymerase inhibitor remdesivir (10–12). Following the same infection protocol described in **Figure 5A**, the compounds were again tested over a range of concentrations in the presence of 1 µM remdesivir. At this concentration, remdesivir alone inhibited viral infection by less than 20%. None of the evaluated nsp13 inhibitors showed significant synergy with remdesivir under these assay conditions (**Supplementary Figure S11B**).

## Discussion

The SARS-CoV-2 protein nsp13 possesses helicase activity and is essential for viral replication and proliferation (24–26, 50). Moreover, nsp13 is the most conserved non-structural protein within the coronavirus family, making it a very promising target for the development of pan-coronavirus antivirals (22). In this study, we developed a robust FRET-based helicase assay and used it for high-throughput inhibitor screening against the SARS-CoV-2 helicase nsp13 testing over 5000 previously characterised pharmaceuticals. We report the identification of FPA-124 and suramin-like compounds as novel inhibitors of nsp13. We performed comparative analyses between the two most promising inhibitors reported in this work (FPA-124 and suramin) and two SARS-CoV-1 nsp13 inhibitors described in the literature (myricetin (39) and SSYA10-001 (37, 38)). We showed that FPA-124 and suramin have lower IC_50_ against nsp13 *in vitro* and lower anti-viral EC_50_ in cell-based assays than the two published compounds.

Colloid formation by aggregation-prone compounds is one of the main sources of false positives in drug discovery (33, 34). Adding non-ionic detergents can reduce colloid formation and thus advise on the specificity of HTS hits (31, 35). All subsequent screens in this series of papers used detergents in the primary screens. Nsp13 inhibition by FPA-124 or suramin showed no or little detergent sensitivity. We performed the SARS-CoV-2 nsp13 HTS in the absence of detergents and obtained 142 primary hits. We then tested a selection of 47 hits with different non-ionic detergents in validation assays. Seven of those tested showed no or only mild changes in IC_50_ values after detergent addition, including FPA-124 and suramin-like compounds (**Supplementary Figure S6-S9**). Myricetin and SSYA10-001, as well as several of the analysed hits from the primary screen, had a significant decrease in potency when assayed in detergent-containing buffers (**Supplementary Figure S8-S9**). Indeed, myricetin has been reported to have an aggregator-like behaviour (51–53) and to form colloidal aggregates detectable by dynamic light scattering (54), raising concerns about the specificity of its reported SARS-CoV-1 nsp13 inhibition (39). We tested a selection of the detergent sensitive hits against a different enzyme, the SARS-CoV-2 RNA-dependent RNA polymerase (RdRp). We found they had similar inhibitory effect on both nsp13 and RdRp further suggesting their mode of action could involve non-specific protein capture by colloid formation. Nonetheless, detergent sensitivity does not completely rule out a *bona fide* inhibitory effect (36). Sometimes this promiscuous activity occurs solely at different concentrations than specific binding-based inhibition. Therefore, some of the compounds that presented a detergent-dependent decrease in inhibition reported in this work may still have *bona fide* inhibitory activity against nsp13 at certain concentrations.

FPA-124 is a cell-permeable Akt inhibitor that induces apoptosis in multiple cancer cell lines (48). We have shown that this compound has an IC_50_ of ∼9 µM and is not detergent-sensitive in *in vitro* assays. FPA-124 was not a hit in any of the other screens in this series (Biochem J, this issue) providing further evidence for specificity. Due to the nature of this compound’s original cellular target, cytotoxicity is expected. Indeed, FPA-124 showed cytotoxicity at 100 µM (**Supplementary Figure S11A**). Suramin, originally synthesised by Bayer in 1916, is a clinically approved drug mainly used to treat river blindness and sleeping sickness (41, 55). We show that suramin and several of its structurally similar compounds are novel inhibitors of SARS-CoV-2 nsp13. Suramin inhibited nsp13 *in vitro* with an IC_50_ of ∼1 µM and inhibited viral growth in cell-based assays with an EC_50_ of ∼10 µM. A wide range of antiviral effects have been reported for suramin, as it inhibits Zika virus, dengue virus, chikungunya virus, HIV, hepatitis C virus, herpes simplex type-1 virus and recently SARS-CoV-2 (41, 56). Suramin seems to inhibit multiple steps in viral infection and replication: it interferes with virus-receptor interaction and hence viral host cell binding and uptake (57, 58), it interferes with viral helicase activities ((45) and this work) and it interferes with viral RNA polymerase activity (46, 59). Suramin is a large symmetrical molecule carrying two polysulfonated naphthyl urea groups containing six negative charges at physiological pH and therefore, the basis for its many targets is likely to be its ability to strongly bind positively charged regions in proteins such as polymerases (42, 44). Indeed, we show that suramin and related compounds also inhibited the SARS-CoV-2 RdRp with micromolar IC_50_s without detergent sensitivity. The polyanionic nature of suramin also confers low cell membrane permeability. However, suramin can be taken up by endocytosis and the uptake rate can be enhanced by liposomal delivery (60).

We validated suramin, suramin-like compounds and FPA-124 as nsp13 inhibitors that could be subjected to careful structural optimization to generate clinically more useful compounds in the hope of increasing antiviral potency and reducing cytotoxicity. Co-structures of nsp13 and these small-molecule inhibitors could highly benefit these optimisation processes. The viral RNA polymerase inhibitor, remdesivir, is currently the only FDA-approved small molecule antiviral for the treatment of COVID-19 patients (10, 61). However, it is far from being a silver bullet as it has been shown to be only modestly effective in treating very sick patients (10, 61). Thus, combining remdesivir with a mechanistically distinct drug (e. g. drugs that target other SARS-CoV-2 enzymatic activities) may improve antiviral efficacy and reduce the likelihood of emerging drug resistance (62). Nonetheless, our experiments in Vero cells showed there was no synergistic inhibition on viral replication when combining remdesivir and our nsp13 inhibitors. Recent studies on host-virus interactions suggest that SARS-CoV-2 nsp13 may also be implicated in immune suppression as it targets several host proteins involved in innate immune signalling pathways such as the interferon pathway and NF-κB pathway (63). Vero E6 cell experiments are useful tools for studying viral replication, but it would be interesting to test our nsp13 inhibitors in other cell lines that have intact innate immune responses in the future (64).

In conclusion, we have identified novel small-molecule inhibitors of SARS-CoV-2 nsp13 helicase. We provide evidence that suramin and FPA-124 can be considered as leads that deserve further evaluation, as both compounds were found to inhibit nsp13 *in vitro* and exhibit antiviral activity against SARS-CoV-2 in a relevant cell culture model.

## Materials and Methods

### Protein expression and purification

FH-nsp13 and GST-nsp13 were expressed in baculovirus-infected insect cells. The coding sequence of SARS-CoV-2 nsp13 (NCBI reference sequence NC_045512.2) was codon-optimised for *S. frugiperda* and synthesized (GeneArt, Thermo Fisher Scientific). Nsp13 was subcloned into the biGBac vector pLIB (65) either to contain an N-terminal 3xFlag-His_6_ tag (sequence: MDYKDHDGDYKDHDIDYKDDDDKGSHHHHHHSAVLQ-nsp13) or to contain an N-terminal GST fusion derived from pGEX-4T-1. Baculoviruses were generated and amplified in Sf9 insect cells (Thermo Fisher Scientific) using the EMBacY baculoviral genome (66). For protein expression Sf9 cells were infected with baculovirus and collected 48 hours post infection, flash-frozen and stored at - 70 °C. FH-nsp13 and GST-nsp13 were affinity purified from insect cell lysates using anti-FLAG M2 Affinity Gel (Sigma-Aldrich) or Glutathione Sepharose 4 Fast Flow resin (GE Healthcare) following the instructions provided by the manufacturer with appropriate modifications. All steps were performed at 4°C. Cell pellets were lysed by Dounce homogenization using a loose-fitting homogenizer pestle in 50 mM HEPES pH 7.4, 200 mM NaCl, 10% Glycerol, 1 mM DTT and the following protease inhibitors: 1 mM PMSF, 0.2 mM AEBSF, 1 μm/mL Aprotinin, 1 µM pepstatin, 10 µM leupeptin, 2 mM Benzamidine, 0.5 µM aprotinin and 1x protease inhibitor cocktail (Complete Ultra Tablets, Roche). The lysate was clarified by centrifugation at 39000 g at 4°C for 60 min. Anti-FLAG M2 resin was added to the FH-nsp13 supernatant and Glutathione Sepharose 4 Fast Flow resin was added to the GST-nsp13 supernatant, and gently rotated for 2 h at 4°C. The resins were transferred into gravity flow columns, washed with wash buffer (50 mM HEPES pH 7.4, 200 mM NaCl, 10% glycerol, 1 mM DTT) and eluted with the respective elution buffer (wash buffer supplemented with 0.1 mg/mL 3xFLAG peptide or wash buffer supplemented with 10 mM reduced glutathione). Both eluates were then fractionated by gel filtration on a Superdex-200 Increase 10/300GL column (GE Healthcare) in buffer containing 50 mM Tris pH 8.0, 200 mM NaCl, 1 mM DTT and 10% glycerol. Fractions containing the desired protein were concentrated, flash frozen in liquid nitrogen and stored at −80°C.

### FRET-based nsp13 helicase assay

A FRET-based fluorescence-quenching approach was designed to monitor nucleic acid strand separation catalysed by nsp13. The assay uses a forked duplex DNA or RNA substrate (15-bp duplex with a 20-nt oligo-dT 5’ overhang). One strand contains a Cy3 fluorophore at the 5’ end (Cy3 strand, DNA or RNA) and the other strand a Black Hole Quencher-2 (BHQ-2, DNA) or an Iowa black RQ quencher (AbRQ, RNA) at the 3’ end (quencher strand). A DNA competitor strand that is complementary to the Cy3 strand prevents substrate reannealing. HPLC-purified DNA and RNA oligonucleotides were purchased from Eurofins genomics and IDT respectively with the following sequences:

**Cy3 strand DNA**: 5’-Cy3-GGTAGTAATCCGCTC-3’

**BHQ-2 strand DNA**: 5’-TTTTTTTTTTTTTTTTTTTTGAGCGGATTACTACC-(BHQ-2)-3’

**Cy3 strand RNA**: 5’-Cy3-GGUAGUAAUCCGCUC-3’

**Iowa Black RQ strand RNA:** 5’-UUUUUUUUUUUUUUUUUUUUGAGCGGAUUACUACC-(AbRQ)-3’

**Competitor strand DNA**: 5’-GAGCGGATTACTACC-3’

The Cy3 strand and the quenching strand were annealed at 1:1.1 ratio at a concentration of 20 µM:22 µM respectively by heating the oligo mix to 75 °C for 5 minutes and gradually cooling it down to 4 °C over 50 minutes. The competitor strand was added at 5 times concentration of the duplex for inhibitor screens and validation assays. A reaction was typically started by adding 10 µl of a 2x substrate solution containing a DNA or RNA substrate and ATP in reaction buffer (20 mM HEPES pH 7.6, 20 mM NaCl, 5 mM MgCl_2_, 1 mM DTT and 0.1 mg/ml BSA) to 10 µl of a 2x enzyme solution containing nsp13 in reaction buffer. Upon unwinding of the duplex by nsp13, Cy3 is no longer quenched by BHQ-2 and thus able to fluoresce. The fluorescence signal was read at 550 nm (excitation) and 570 nm (emission) with 5 nm bandwidth using a Tecan Infinite M1000, or at 545 nm (excitation) and 575 nm (emission) with 10 nm bandwidth using a Tecan Spark microplate reader.

### Radiolabelled-based nsp13 helicase assay

The oligonucleotides used in the radiolabelled assays are the following:

**Short strand DNA**: 5’-GGTAGTAATCCGCTC-3’

**5’ overhang strand DNA**: 5’-TTTTTTTTTTTTTTTTTTTTGAGCGGATTACTACC-3’

**Short strand RNA**: 5’-GGUAGUAAUCCGCUC-3’

**5’ overhang strand RNA:** 5’-UUUUUUUUUUUUUUUUUUUUGAGCGGAUUACUACC-3’

**Competitor strand DNA**: 5’-TTTTTTTTTTTTTTTTTTTTGAGCGGATTACTACC-3’

DNA or RNA oligonucleotide containing the 5’ overhang (**5’ overhang strand**) was radiolabelled at the 5’ ends with [γ-^32^P] ATP by T4 PNK. The unmodified DNA version of the 5’ overhang strand was used as the competitor strand. The unwinding reactions were carried out at room temperature in helicase buffer (20 mM HEPES pH 7.5, 20 mM NaCl, 5 mM MgCl_2_, 100 /mL BSA, 1 mM DTT, RNase inhibitor Murine (NEB) 2 units/µl) containing 1mM ATP for 50 nM DNA substrate or 2 mM ATP for 100 nM RNA substrate. The unwinding reactions were quenched at various times with a quench solution containing 50 mM EDTA. The aliquots then were mixed with 5x loading buffer (TBE buffer, 15% Ficoll, 1 µg/ml bromophenol blue, 1 µg/ml xylene cyanol FF). The products of the unwinding reactions were separated on a 20% native-PAGE (0.5X TBE). The gels were dried and then exposed to a Phospho-Imager screen overnight, imaged using Phospho-Imager software.

### Nsp13 High-Throughput Screening Assay

The screen was performed in 384-well Greiner black flat-bottom plates (Greiner 781076). A custom compound library containing over 5000 compounds assembled from commercial sources (Sigma, Selleck, Enzo, Tocris, Calbiochem, and Symansis) were distributed in 24 plates using an Echo 550 (Labcyte) across column 3 to column 22. 2.5, or 12.5 nL of a 10 mM stock of the compounds dissolved in DMSO were arrayed and pre-dispensed into the assay plates using an Echo 550 (Labcyte), before being sealed and stored at -80 °C until screening day. All 384 wells on the plates contain 1 µL DMSO. First, 10 µL 2x Nsp13 mix (6 nM FH-nsp13) in HTS assay buffer (20 mM HEPES pH 7.6, 20 mM NaCl, 5 mM MgCl_2_, 1 mM DTT and 0.1 mg/ml BSA) was dispensed into columns 2 - 23 of the plates to incubate with the compounds for 10 min at room temperature. Then 10 µL 2x substrate mix (200 µM ATP, 360 nM DNA substrate, 1800 nM competitor) in HTS assay buffer was dispensed to start the enzymatic reaction. The plates were then spun briefly and transferred to a Tecan microplate reader to monitor changes in fluorescence signal in a kinetic mode. The fluorescence signal was first read at 2 min of reaction and then read every 1.5 min for 10 cycles. The screening for nsp13 inhibitors was done twice, one with a final compound concentration of 1.25 µM and one with 6.25 µM.

### Data analysis

The first fluorescence signal at 2 min after reaction start in the screen was used as the initial velocity (V_0_) to compare nsp13 activity in different wells. MATLAB was used to process data. The initial velocity for each compound is first normalised against the DMSO controls in column 23 of each plate as following:

***Normalised_V_0_ = [V_0__of_compound – mean(no_enzyme)]/[mean(V_0__DMSO) – mean (no_enzyme)]***

Because there was a time delay in reading different wells by the microplate reader, reactions in wells that were read later progressed further than wells that were read earlier in each cycle, resulting in higher signals towards the end of a microplate. This positional variation is corrected as following: For each plate, a linear regression of Normalised_V_0_ is fitted against well numbers of wells in between column 2 and column 23. A slope value (S) and an intercept value (I) from this linear regression were obtained and used to do the positional correction for the N_th_ well of a 384-well plate.

***Corrected_V_0_= Normalised_V_0_/(S*N+I)***

For endpoint analysis, fluorescent signals from 6.5 min, 8 min and 9.5 min were taken average when the enzymatic reaction had finished in the DMSO control wells. Compounds that gave more than 10% inhibition in endpoint analysis were considered hits and were collated with hits from initial velocity analysis. Plate 14 from the 1.25 µM screen, plate 1 and 5 from 6.25 µM screen were excluded from data analysis because of faulty handling of the plates. In total, there were 339 hits after initial selection.

For preliminary evaluation of the 339 hits, a kinetic plot in which the y-axis shows fluorescent signals, and the x-axis shows timepoints was drawn for each hit. In addition, signals from 10 wells that were before and after the hit well in a 384-well plate were plotted in the same graph with the hit. Because the wells were close to each other, their reaction time was similar and thus one would expect their curves overlap if there was no inhibition of the reaction. And a real hit would produce lower signal than the neighbouring wells. We manually inspected the curves for all the 339 hits and confirmed that 142 of them gave obvious lower signal than their neighbouring wells.

To be used as a reference for the hit selection method described above, the Z-score was calculated for each compound (Supplementary Figure S3G) as following:

***Z-score=[Corrected_V_0__of_a_compound – mean(Corrected_V_0_)]/SD(Corrected_V_0_),***

where mean(Corrected_V0) is the average of corrected initial velocity values of all compounds and SD(Corrected_V0) is the standard deviation of corrected initial velocity values of all compounds.

To assess the quality of the screen, the Z-factor was calculated for each plate based on corrected initial velocity values as following:

***Z-factor=1–[3*SD(Corrected_V_0__of_DMSO) 3*SD(no_enzyme)] / [mean(Corrected_V_0__of_DMSO) – mean(no_enzyme)]***

The average Z-factor for the screen after positional correction is 0.53. This value suggests the screen is good, as described by Zhang et al. (67).

### Tested Drugs

SSYA001-10, myricetin and remdesivir together with the compounds selected for *in vitro* validation and cell-based studies were purchased and resuspended following manufacturer’s instructions. See **Supplementary Table 4** for more details.

### Gel-based helicase assays

A compound volume of 0.5 µl at a concentration of 300 µM was incubated with 5 µl protein mix containing 4 nM FH-nsp13 for 10 min at room temperature. The reaction was started by adding 5 µl substrate mix containing 100 nM forked RNA substrate (Cy3 RNA strand and unlabelled 5’ overhang RNA strand) and 4 mM ATP in a reaction buffer containing 20 mM HEPES pH 7.6, 5 mM MgCl_2_, 20 mM NaCl, 1 mM DTT, 0.1 mg/ml BSA and 0.01% Triton X-100. The reaction mix also contained 1 µM unlabelled 15-mer RNA as a competitor to trap the displaced overhang strand. After 5 minutes at room temperature, the reaction was quenched by adding 5 µl quenching buffer (20 mM EDTA, 0.5% SDS, 2 mg/ml proteinase K, 10% glycerol) and incubating at 37 °C for 5 min. Reaction products were then analysed by running on a 20% non-denaturing PAGE gel in 0.5X TBE buffer (Life Technologies) for 20 min at room temperature and 200 V. An Amersham Imager 600 was used to image Cy3 fluorescence.

### SARS-CoV-2 production

Fifty percent confluent monolayers of Vero E6 cells (courtesy of National Institute for Biological Standards and Control, NIBSC) were infected with the SARS CoV-2 strain England/2/2020 (courtesy of Public Health England, PHE) at an MOI of approx. 0.001. Cells were washed once with DMEM (Sigma; D6429), then 5 ml virus inoculum made up in DMEM was added to each T175 flask and incubated at room temperature for 30 minutes. DMEM + 1% FCS (Biosera; FB-1001/500) was added to each flask. Cells were incubated at 37°C, 5% CO2 for 4 days until extensive cytopathogenic effect was observed. Supernatant was harvested and clarified by centrifugation at 2000 rpm for 10 minutes in a benchtop centrifuge. Supernatant was aliquoted and frozen at -80°C.

### Recombinant mAb production

Heavy and light chain variable regions for CR3009, were synthesized (Genewiz) based on the GenBank sequences with regions of overlap to restriction digested human IgG1 vectors for assembly cloning (NEB) to produce plasmids: CR3009HC and CR3001KC. N-protein specific mAb CR3009 was produced by co-transfecting Expi293F cells (Life Technologies) in suspension growing at 37oC in 8% CO2 atmosphere in FreeStyle 293T medium (Life Technologies) with the plasmids. The supernatants were harvested 6-8 days post-transfection. as per the original study describing this mAb (68). The CR3009 Mab were purified by affinity chromatography using a 5 ml Protein G column (Cytiva) attached to an AKTA Pure system. Upon loading, the column was washed with PBS and bound mAbs were eluted with 0.1M glycine pH 2.2 and immediately neutralized with 1M Tris, pH 8.0. The mAb-containing fractions were pooled and subjected to size exclusion chromatography using a Superdex200 16/600 prep grade column. The purified mAb CR3009 was labelled with Alexa Fluor 488-NHS (Cat#1812 Jena Biosciences) according to the instructions from the manufacturer.

### Viral inhibition assay

1.5*10^3 Vero E6 cells (NIBC, UK) resuspended in DMEM containing 10% FBS were seeded into each well of 96-well imaging plates (Greiner 655090) and cultured overnight at 37C and 5% CO2. The next day, a 5x solution of compounds were generated by dispensing 10mM stocks of compounds into a v-bottom 96-well plate (Thermo 249946) and back filling with DMSO to equalise the DMSO concentration in all wells using an Echo 550 (Labcyte) before resuspending in DMEM containing 10% FBS. The assay plates with seeded VERO cells had the media replaced with 60µl of fresh growth media, then 20µl of the 5x compounds were stamped into the wells of the assay plates using a Biomek Fx automated liquid handler. Finally, the cells were infected by adding 20µl of SARS-CoV-2 with a final MOI of 0.5 PFU/cell. 22h post infection, cells were fixed, permeabilised, and stained for SARS-CoV-2 N protein using Alexa488-labelled-CR3009 antibody produced in-house (see section for Recombinant mAb production) and cellular DNA using DRAQ7 (Abcam). Whole-well imaging at 5x was carried out using an Opera Phenix (Perkin Elmer) and fluorescent areas and intensity calculated using the Phenix-associated software Harmony (Perkin Elmer).

### CRediT Contribution

**Jingkun Zeng**: Conceptualization, Methodology, Investigation, Software, Formal analysis, Data Curation, Validation, Visualization, Writing – original draft, Writing - review & editing. **Florian Weissmann**: Conceptualization, Methodology, Investigation, Formal analysis, Validation, Visualization, Writing – original draft, Writing - review & editing. **Agustina P. Bertolin**: Conceptualization, Methodology, Investigation, Formal analysis, Validation, Visualization, Writing – original draft, Writing - review & editing. **Viktor Posse**: Conceptualization, Methodology, Investigation, Writing - review & editing. **Berta Canal**: Conceptualization, Investigation, Writing - review & editing. **Rachel Ulferts**: Investigation. **Mary Wu**: Investigation. **Ruth Harvey**: Resources. **Saira Hussain**: Resources. **Jennifer Milligan**: Investigation. **Svend Kjaer**: Resources, Supervision. **Chloe Roustan**: Resources. **Annabel Borg**: Resources. **Laura McCoy**: Resources **Lucy S. Drury**: Resources. **John McCauley**: Resources, Supervision. **Michael Howell**: Resources, Supervision. **Rupert Beale**: Resources, Supervision. **John F.X Diffley**: Conceptualization, Methodology, Formal analysis, Project administration, Writing - review & editing, Funding acquisition, Supervision.

## Acknowledgements

We thank Anne Early for assistance. This work was supported by the Francis Crick Institute, which receives its core funding from Cancer Research UK (FC001066, FC001030), the UK Medical Research Council (, FC001030), and the Wellcome Trust (FC001066, FC001030). This work was also funded by a Wellcome Trust Senior Investigator Award (106252/Z/14/Z) to J.F.X.D. FW and BC have received funding from the European Union’s Horizon 2020 research and innovation programme under the Marie Skłodowska-Curie grant agreement Nos 844211 and 895786. JZ has received funding from a Ph.D. fellowship awarded by Boehringer Ingelheim Fonds.

---

**Supplementary Figure S1.**
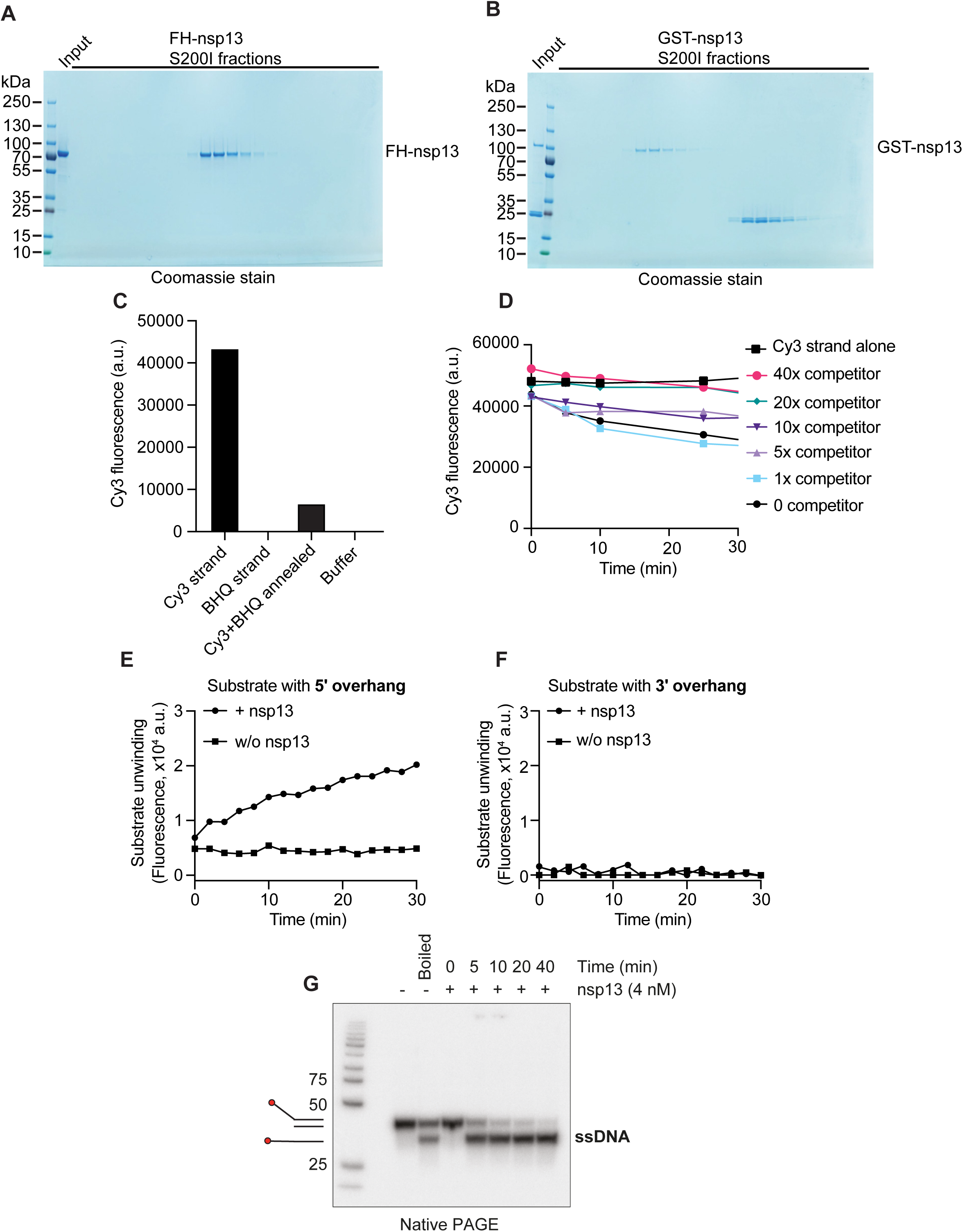
SARS-CoV-2 nsp13 protein expression and FRET-based helicase assay development and optimization. (**A**, **B**) FH-nsp13 and GST-nsp13 were purified by gel filtration on a Superdex 200 Increase 10/300 GL column (S200I) and fractions were analysed by SDS-PAGE and Coomassie staining. (**C**) Emission of fluorescence by 50 nM Cy3 strand could be efficiently quenched by 55 nM BHQ-2 strand after the annealing protocol. (**D**) Annealing dynamics of the Cy3 strand and BHQ strand at room temperature. Without the competitor strand (0 competitor), 50 nM Cy3 strand slowly annealed with 50 nM BHQ-2 strand resulting in quenching of fluorescence by approximately 40% over 30 min. Adding increasing fold of the competitor strand (relative to 50 nM) results in less quenching by disrupting spontaneous annealing between the Cy3 and BHQ strands. (**E**, **F**) Unwinding of 50 nM DNA duplex by 1 nM FH-nsp13 was observed with a duplex with 5’ dT overhang but not with a duplex with 3’ dT overhang. (**G**) A native gel-based assay was used to visualise unwinding of DNA duplexes by nsp13. The 5’ end of the longer strand of the DNA duplex was radiolabelled. Strand separation of the duplex by nsp13 was observed over time.

**Supplementary Figure S2.**
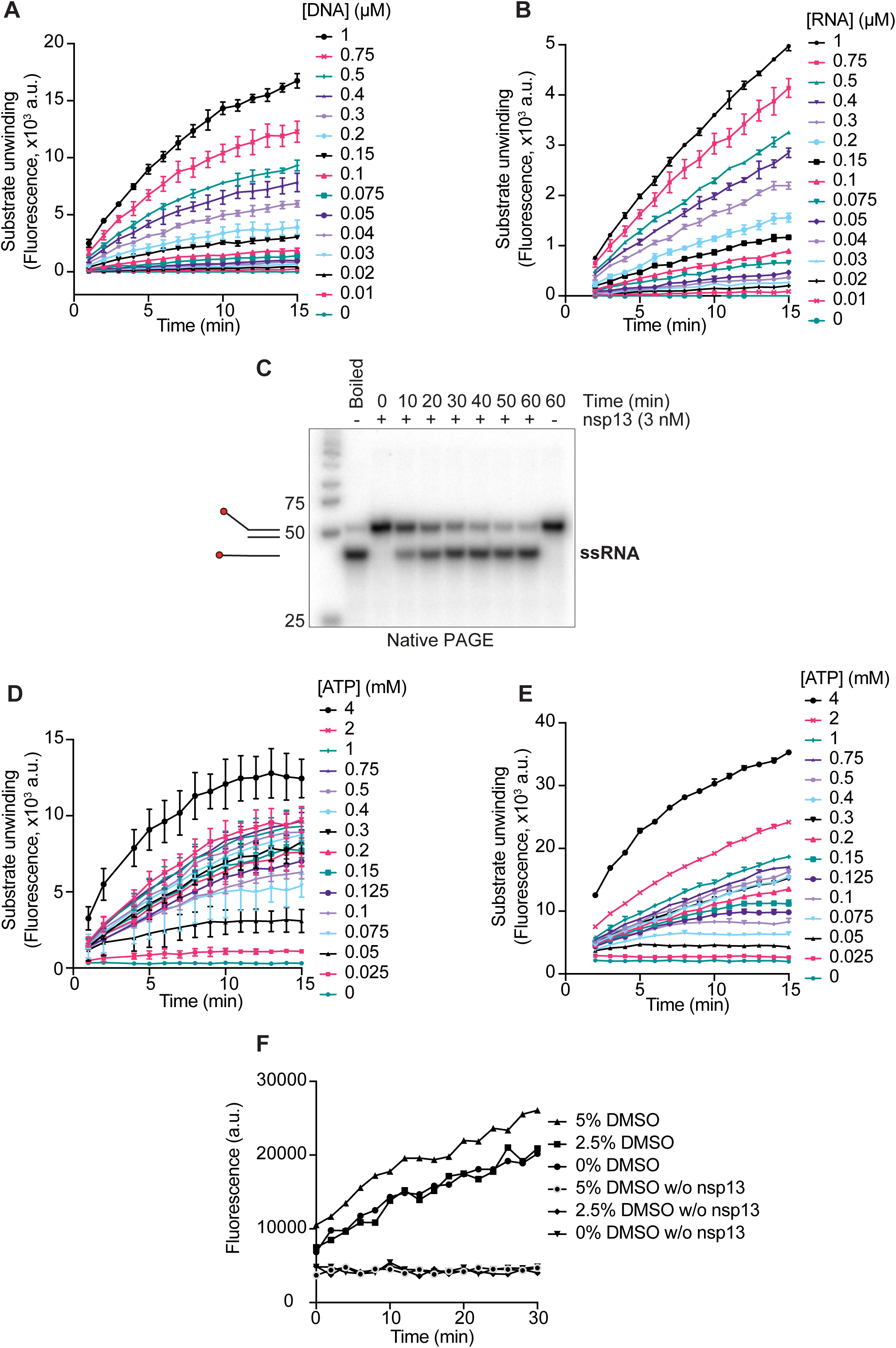
SARS-CoV-2 nsp13 enzyme kinetics. (**A-B,D-E**) Nsp13 activity was tested with increasing concentrations of ATP or the nucleic acid substrates to determine the Michaelis constants (K_m_) shown in Figure 2. (**A**) DNA substrate titration. Conditions: 1 nM FH-nsp13, 1 mM ATP, indicated DNA substrate concentrations. (**B**) RNA substrate titration. Conditions: 1 nM FH-nsp13, 1 mM ATP, indicated RNA substrate concentrations. (**C**) A native gel-based assay was used to visualise unwinding of RNA duplexes by FH-nsp13. The 5’ end of the longer strand of the RNA duplex was radiolabelled. Strand separation of the duplex by nsp13 was observed over time. (**D**) ATP titration in the presence of a DNA substrate. Conditions: 2 nM FH-nsp13, 200 nM DNA substrate, indicated ATP concentrations. (**E**) ATP titration in the presence of the RNA substrate. Conditions: 3 nM FH-nsp13, 200 nM RNA substrate, indicated ATP concentrations. (**F**) Addition of up to 5% DMSO did not disrupt nsp13 activity or annealing properties of nucleic acid substrates.

**Supplementary Figure S3.**
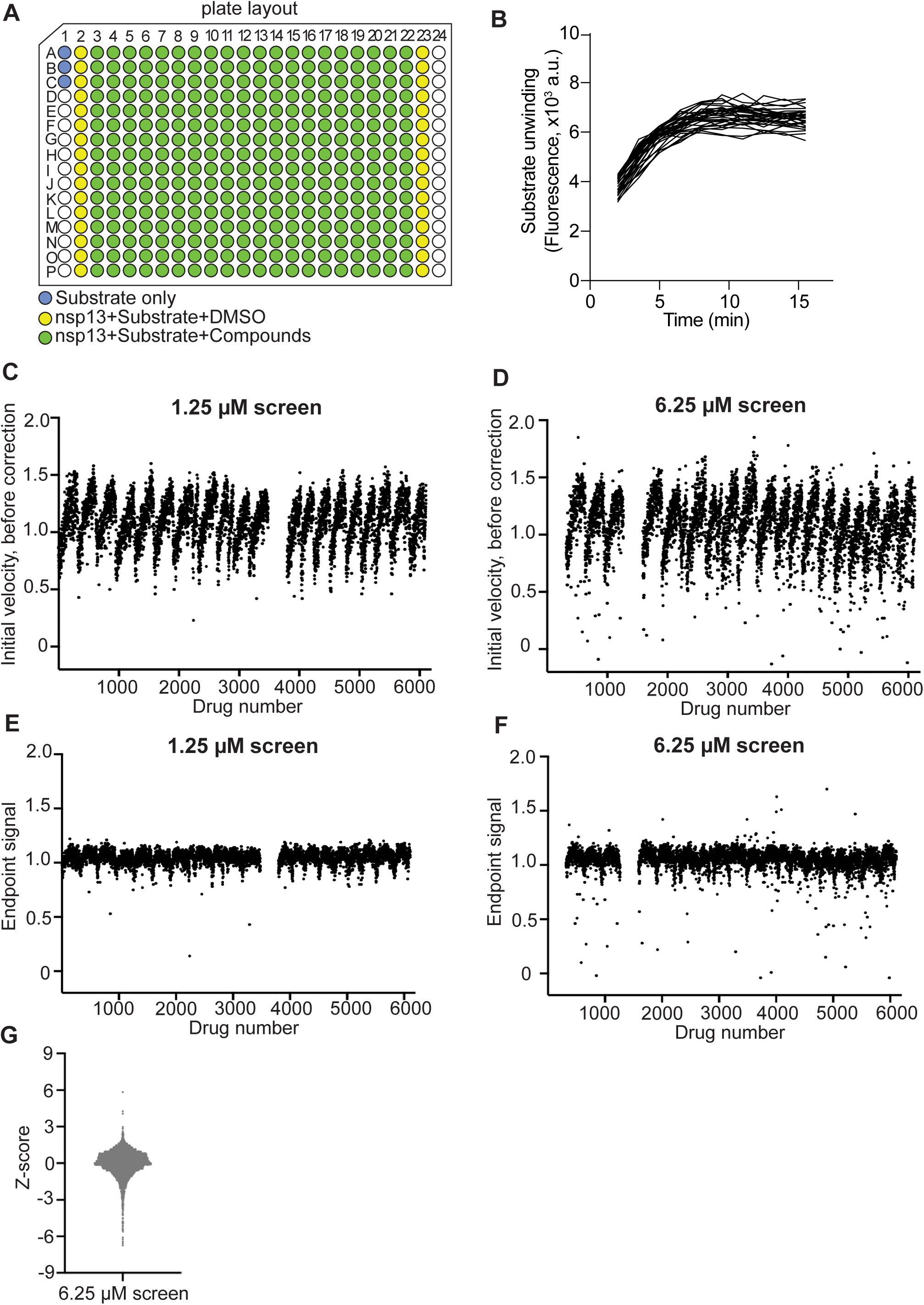
High-throughput SARS-CoV-2 nsp13 inhibitor screen. (**A**) Layout of 384-well plates used in the HTS. (**B**) Representative time-course data showing nsp13 DNA duplex unwinding activity in a 384-well plate format as used in the HTS. (**C, D**) Normalised initial velocity values for all the compounds in the HTS, before positional correction. (**E, F**) Normalised endpoint signals for all the compounds in the HTS. (**G**) Distribution of Z-scores of all the compounds tested in the 6.25 µM screen.

**Supplementary Figure S4.**
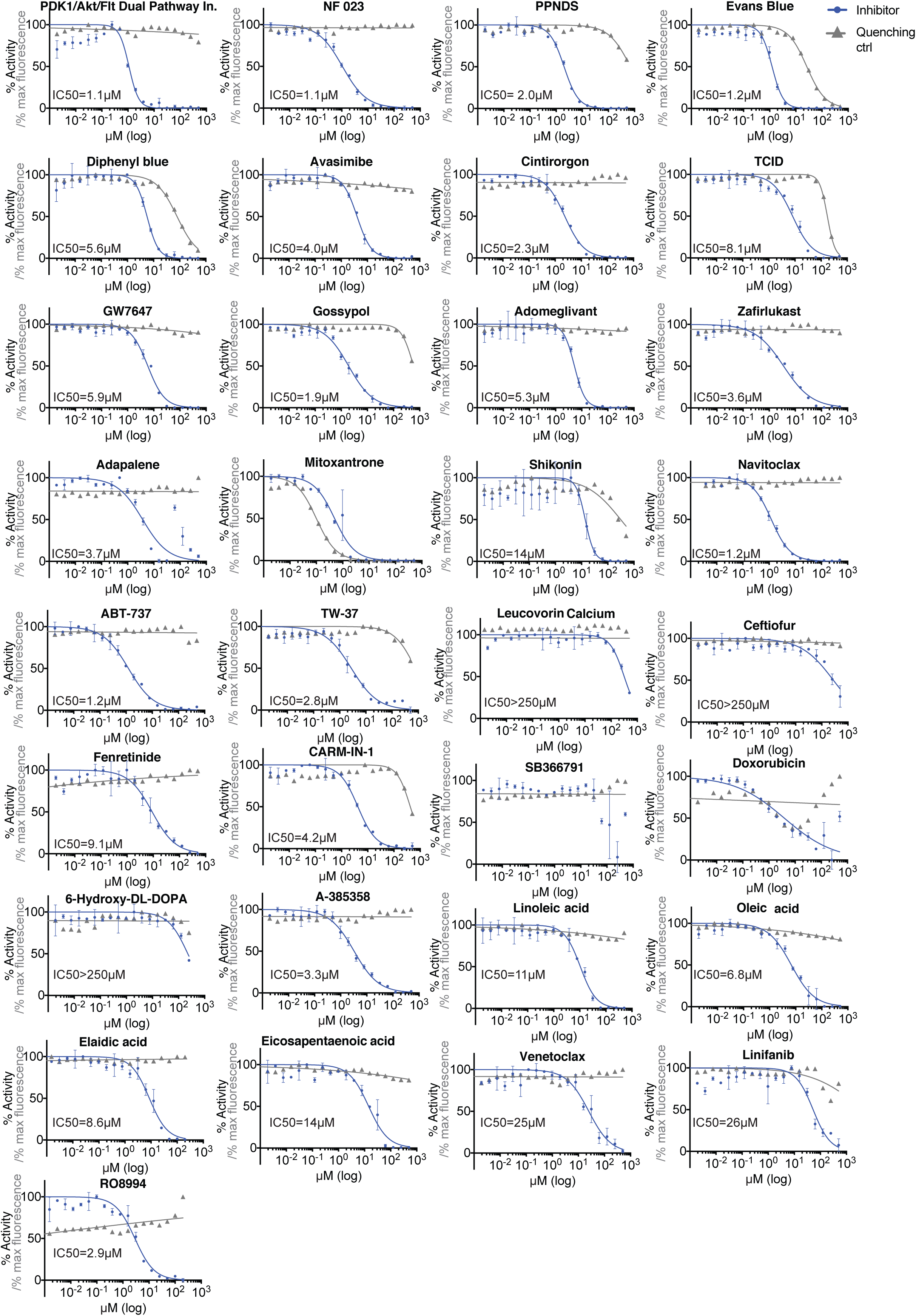
Concentration-response curves for validation of selected hits identified in the SARS-CoV-2 nsp13 HTS. The experiment (blue) was performed with 180 nM fluorogenic DNA substrate, 0.1 mM ATP, 1.5 nM nsp13 and in the absence of detergent. Fluorescence quenching properties of the compounds were tested at the same compound concentrations to rule out false positives that interfered with fluorescence emission (grey). Fluorescence intensities were measured using 180 nM unannealed Cy3 strand in the absence of a quencher strand.

**Supplementary Figure S5.**
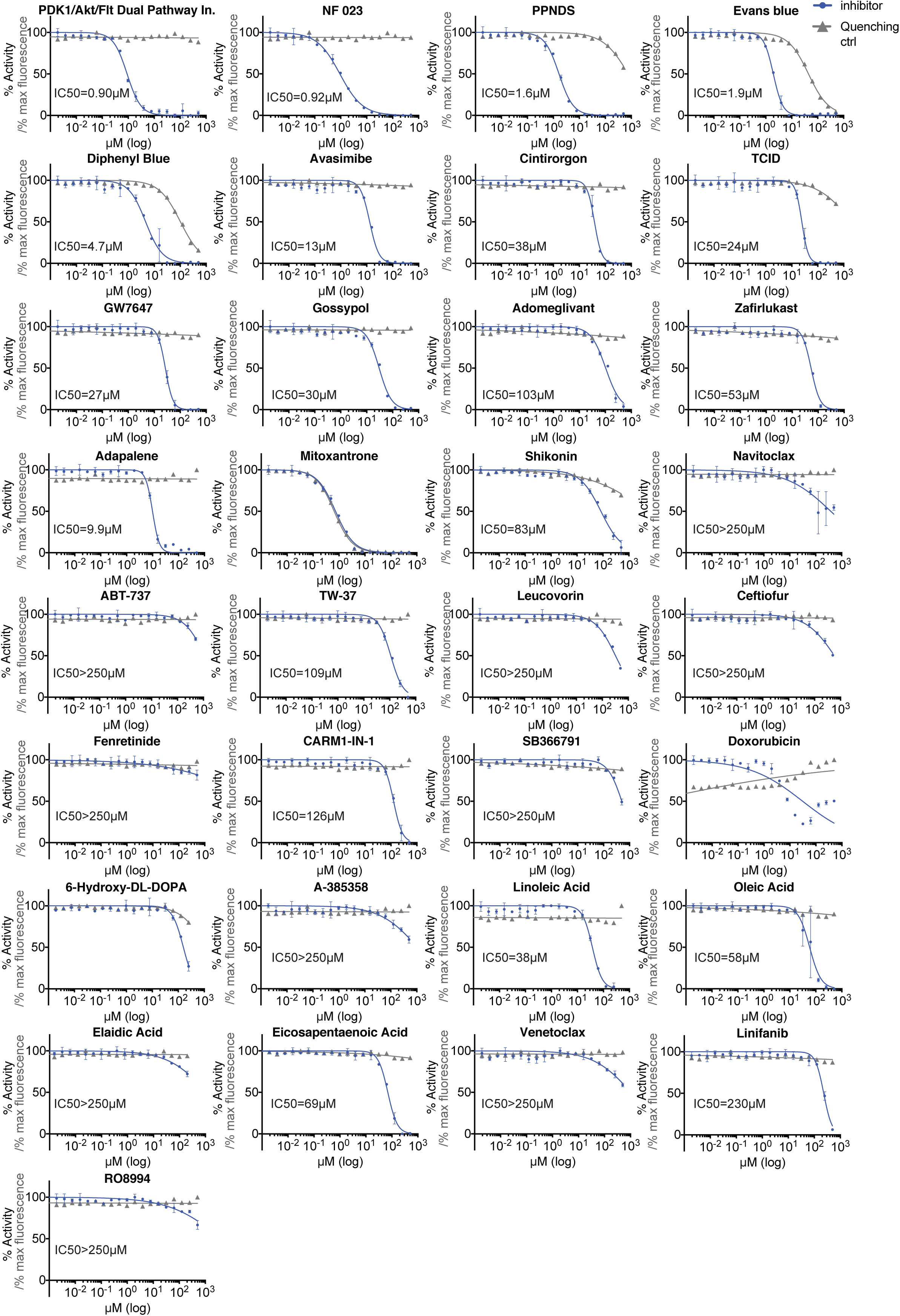
Concentration-response curves for validation of selected hits identified in SARS-CoV-2 nsp13 HTS. The experiment (blue) was performed using 180 nM fluorogenic RNA substrate, 2 mM ATP, 1 nM nsp13 and in the presence of 0.02% Tween-20. Fluorescence quenching properties of the compounds were also tested (grey) to rule out false positives that interfered with fluorescence emission.

**Supplementary Figure S6.**
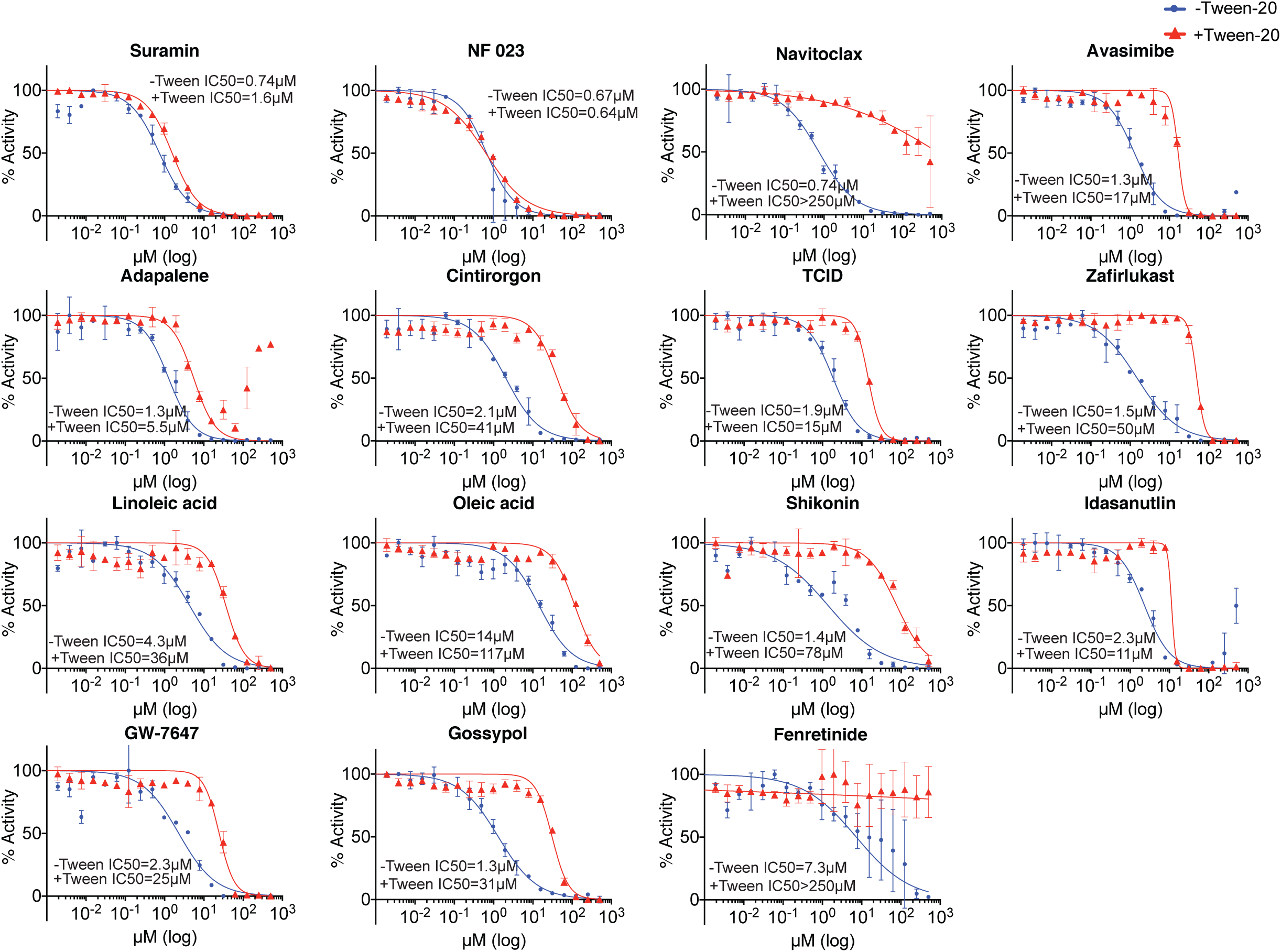
Concentration-response curves for validation of selected hits identified in SARS-CoV-2 nsp13 HTS. The experiment was performed using 180 nM fluorogenic DNA substrate, 0.1 mM ATP, 0.5 nM nsp13 in the presence (red) or absence (blue) of 0.02% Tween-20 in the reaction.

**Supplementary Figure S7.**
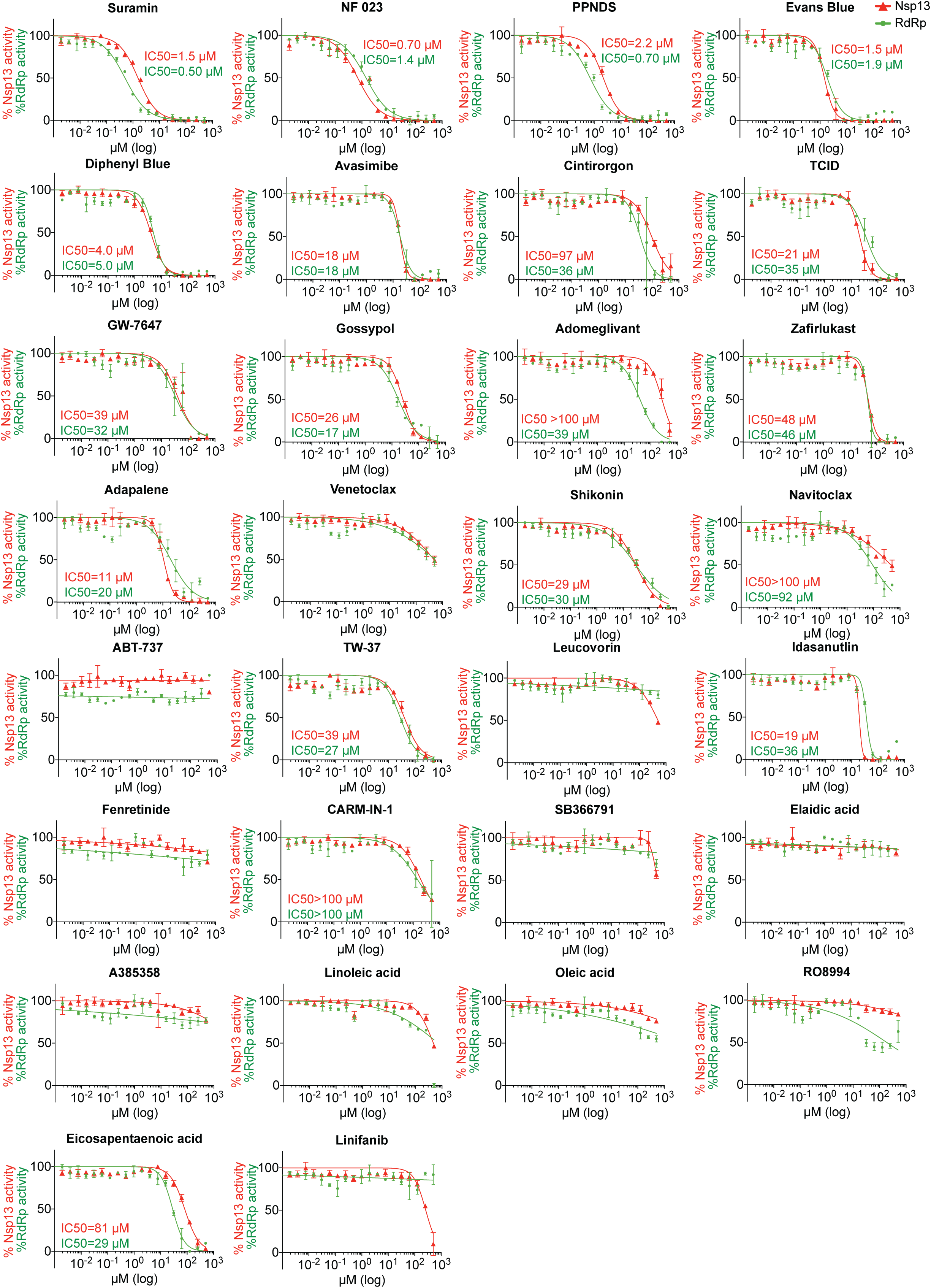
Concentration-response curves of SARS-CoV-2 nsp13 and RdRp activity for selected compounds. Nsp13 helicase activity was assessed using the helicase assay described in this manuscript and was performed with 100 nM fluorogenic RNA substrate, 0.3 mM ATP, 3 nM nsp13 in RdRp assay buffer (see below) with 0.02% Tween-20. RdRp activity was measured using the strand displacement (SD) assay described in detail in Bertolin et al (Biochem J, this issue) and was performed with 100 nM of fluorogenic RNA substrate, 100 nM of RdRp complex (Sf nsp12-3F/7L8 preincubated with Sf 7H8 at a 1:3 ratio) in RdRp assay buffer (20 mM HEPES pH 7.5, 10 mM KCl, 1 mM DTT, 5 mM MgCl_2_, 2 mM MnCl_2_, 0.1 mg/ml BSA) with 0.02% Tween-20. Nsp13 activity in the helicase assay is depicted in red and RdRp activity in the SD assay is depicted in green.

**Supplementary Figure S8.**
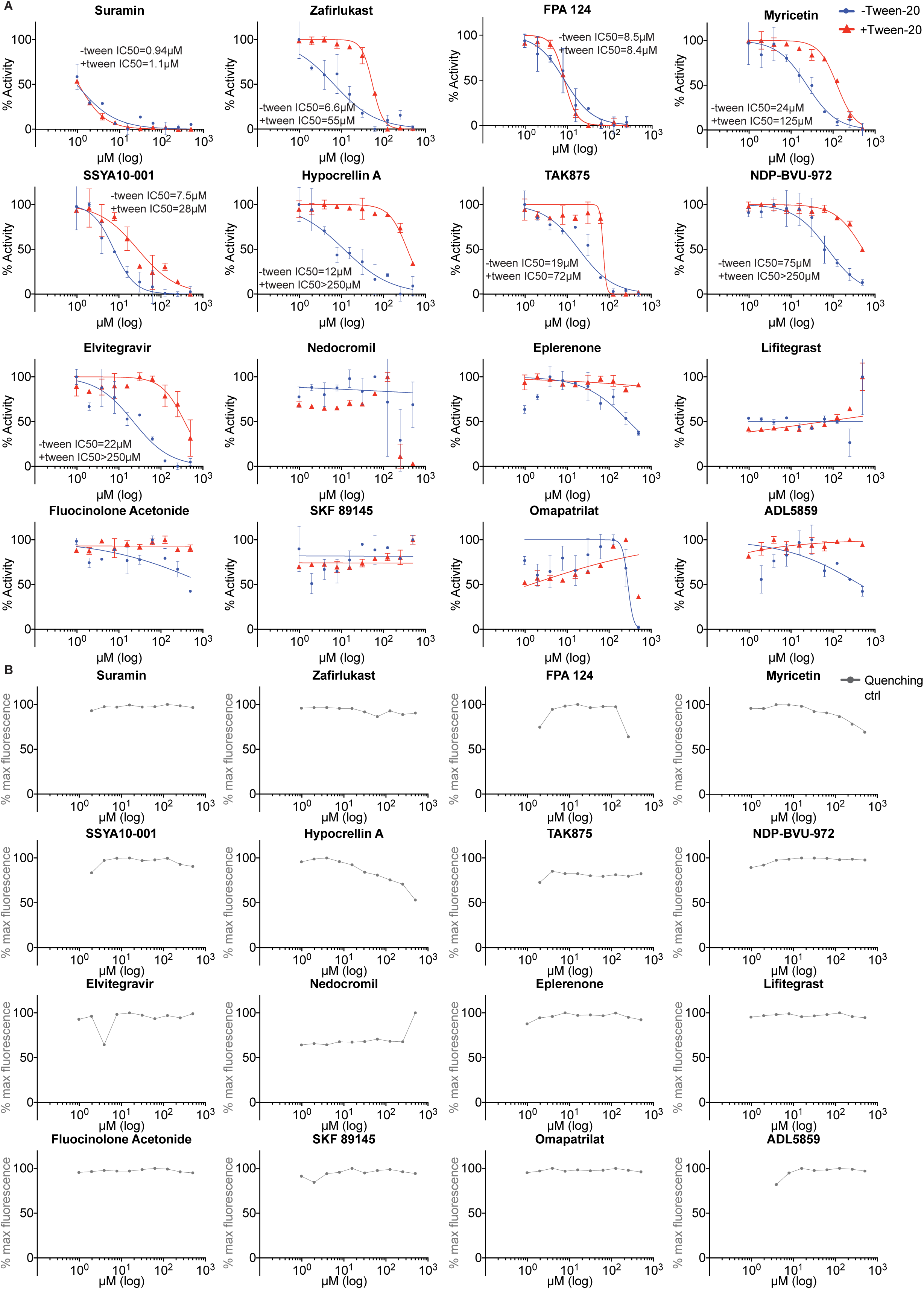
Concentration-response curves for validation of selected hits identified in SARS-CoV-2 nsp13 HTS. (**A**) The experiment was performed with 180 nM fluorogenic RNA substrate, 0.1 mM ATP, 1 nM nsp13 in the presence (red) or absence (blue) of 0.02% Tween-20. (**B**) Fluorescence quenching properties of the compounds were also tested (grey) to rule out false positives that interfered with fluorescence emission.

**Supplementary Figure S9.**
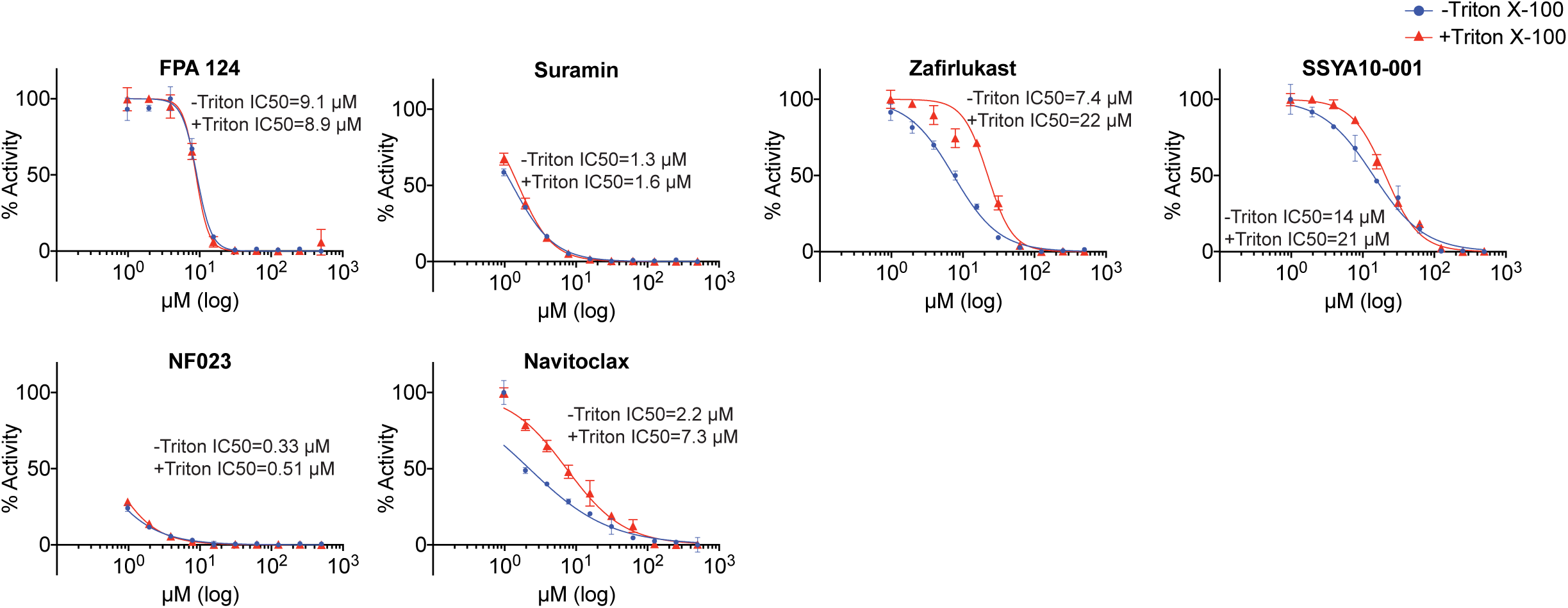
Concentration-response curves for validation of selected hits identified in SARS-CoV-2 nsp13 HTS. The experiment was performed using 180 nM fluorogenic RNA substrate, 0.1 mM ATP, 1 nM nsp13 in the presence (red) or absence (blue) of 0.01% Triton X-100.

**Supplementary Figure S10.**
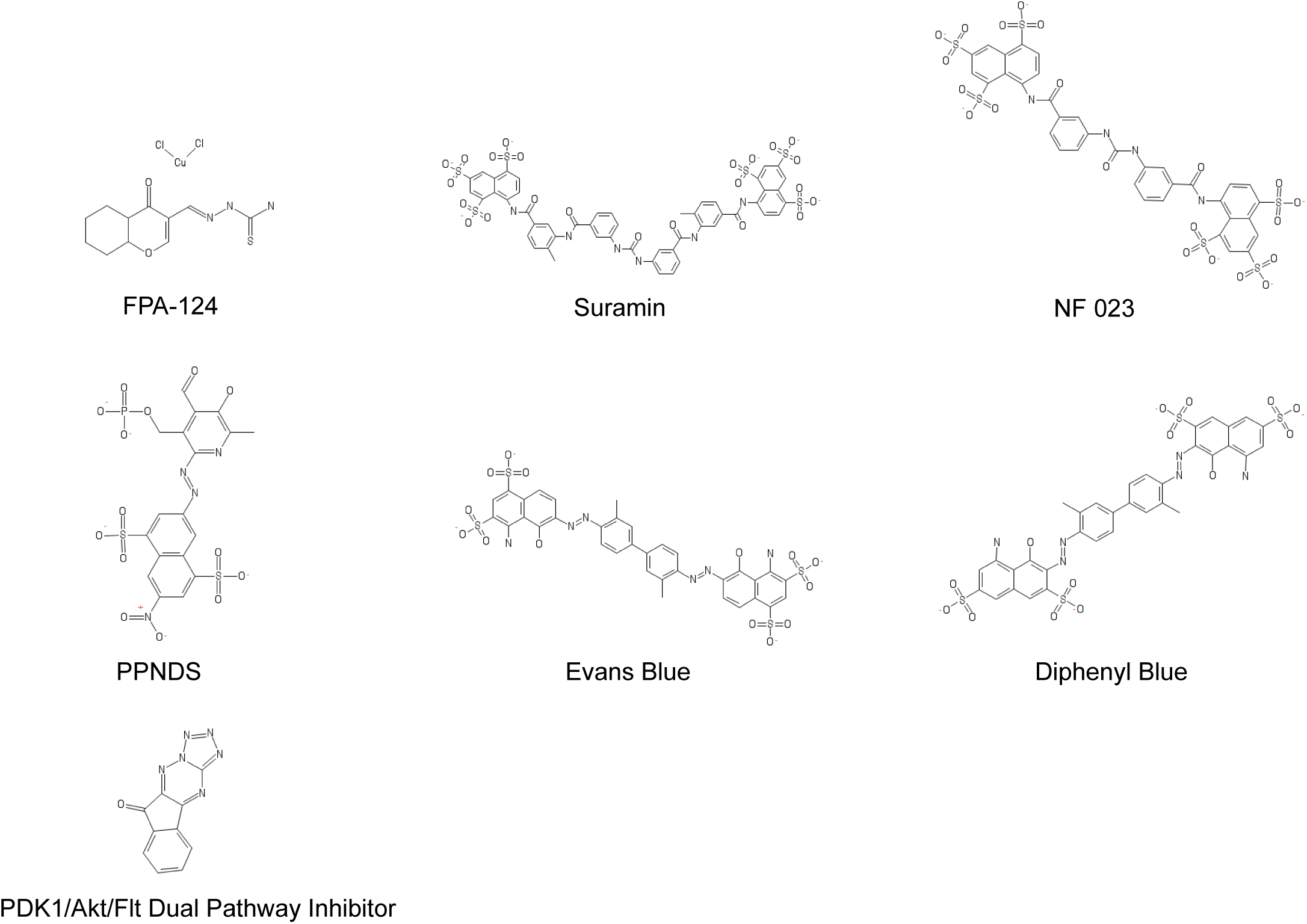
Chemical structures of selected nsp13 inhibitors. FPA-124 data is shown in Table 2. Suramin, NF 023, PPNDS, Evans Blue, Diphenyl Blue, PDK1/Akt/Flt Dual Pathway Inhibitor data are shown in Table 1.

**Supplementary Figure S11.**
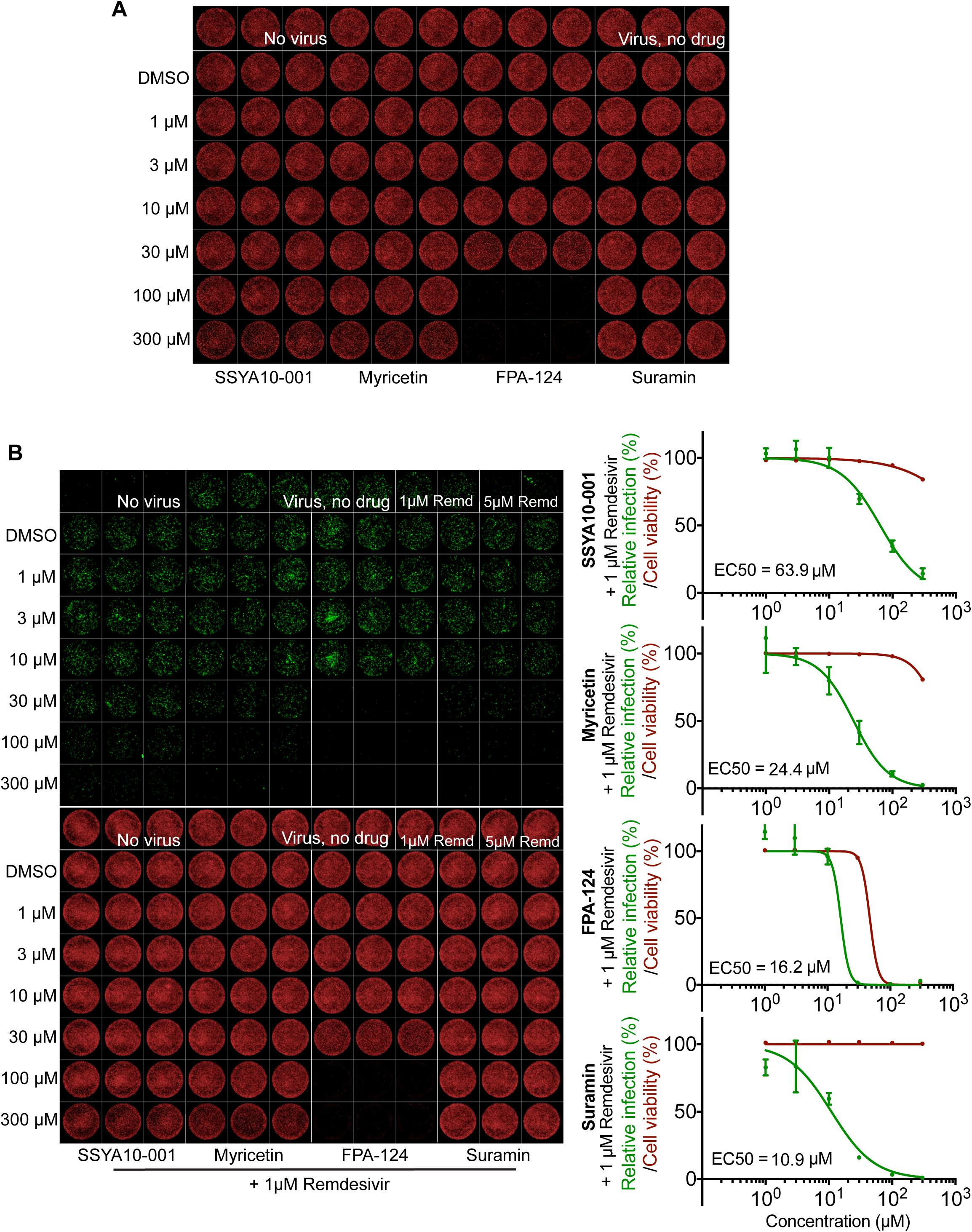
Cytotoxicity analysis and comparative dose–response curves of selected antiviral compounds against SARS-CoV-2 in cell culture. (**A**) SSYA10-001, myricetin, suramin and FPA-124 cytotoxicity analysis of DRAQ7-stained Vero E6 cell. Representative images of one experiment in triplicate shown. (**B**) Anti-SARS-CoV-2 activities of SSYA10-001, myricetin, suramin and FPA-124 in combination with 1 µM remdesivir. Left, representative images showing N protein immunofluorescence (green) and DRAQ7 staining (red) of one experiment in triplicate shown. Right, dose-response curve analysis. Viral infection was calculated as the area of viral plaques stained for N protein and cell viability as the area of cells stained for DRAQ7. Data is plotted as percentage compared to 1 µM remdesivir only treated wells (100%). Values represent mean and standard deviation (SD) of three replicates. FIJI software was used to calculate areas and Prism software to calculate EC_50_ values.

**Supplementary Table 1.** Inhibitory activity of 30 selected compounds against SARS-CoV-2 nsp13 and RdRp holoenzyme

| Compound name | IC<sub>50</sub> (μM) |  |
| --- | --- | --- |
|  | nsp13<sup>a</sup> | RdRp<sup>b</sup> |
| NF 023 | 0.7 | 1.4 |
| Suramin | 1.5 | 0.5 |
| PPNDS | 2.2 | 0.7 |
| Evans Blue | 1.5 | 1.9 |
| Diphenyl Blue | 4.0 | 5.0 |
| Adapalene | 11 | 20 |
| Avasimibe | 18 | 18 |
| Idasanutlin | 19 | 36 |
| TCID | 21 | 35 |
| Gossypol | 26 | 17 |
| Shikonin | 29 | 30 |
| TW-37 | 39 | 27 |
| GW7647 | 39 | 32 |
| Zafirlukast | 48 | 46 |
| Eicosapentaenoic acid | 81 | 29 |
| Cintirorgon | 97 | 36 |
| Adomeglivant | >100 | 39 |
| Navitoclax | >100 | 92 |
| CARM1-IN-1 | >100 | >100 |
| Linifanib/ABT-869 | No inhibition | No inhibition |
| ABT-737 | No inhibition | No inhibition |
| Linoleic acid | No inhibition | No inhibition |
| Oleic acid | No inhibition | No inhibition |
| RO8994 | No inhibition | No inhibition |
| A-385358 | No inhibition | No inhibition |
| Fenretinide | No inhibition | No inhibition |
| Elaidic acid | No inhibition | No inhibition |
| Venetoclax/ABT-199 | No inhibition | No inhibition |
| Leucovorin Calcium | No inhibition | No inhibition |
| SB-366791 | No inhibition | No inhibition |

<sup>a</sup> Experimental conditions: RNA (nsp13 substrate), 0.3 mM ATP, 0.02% Tween-20 in RdRp assay buffer
<sup>b</sup> Experimental conditions: RNA (RdRp substrate), 0.3 mM each NTP, 0.02% Tween-20 in RdRp assay buffer

**Supplementary Table 2.**
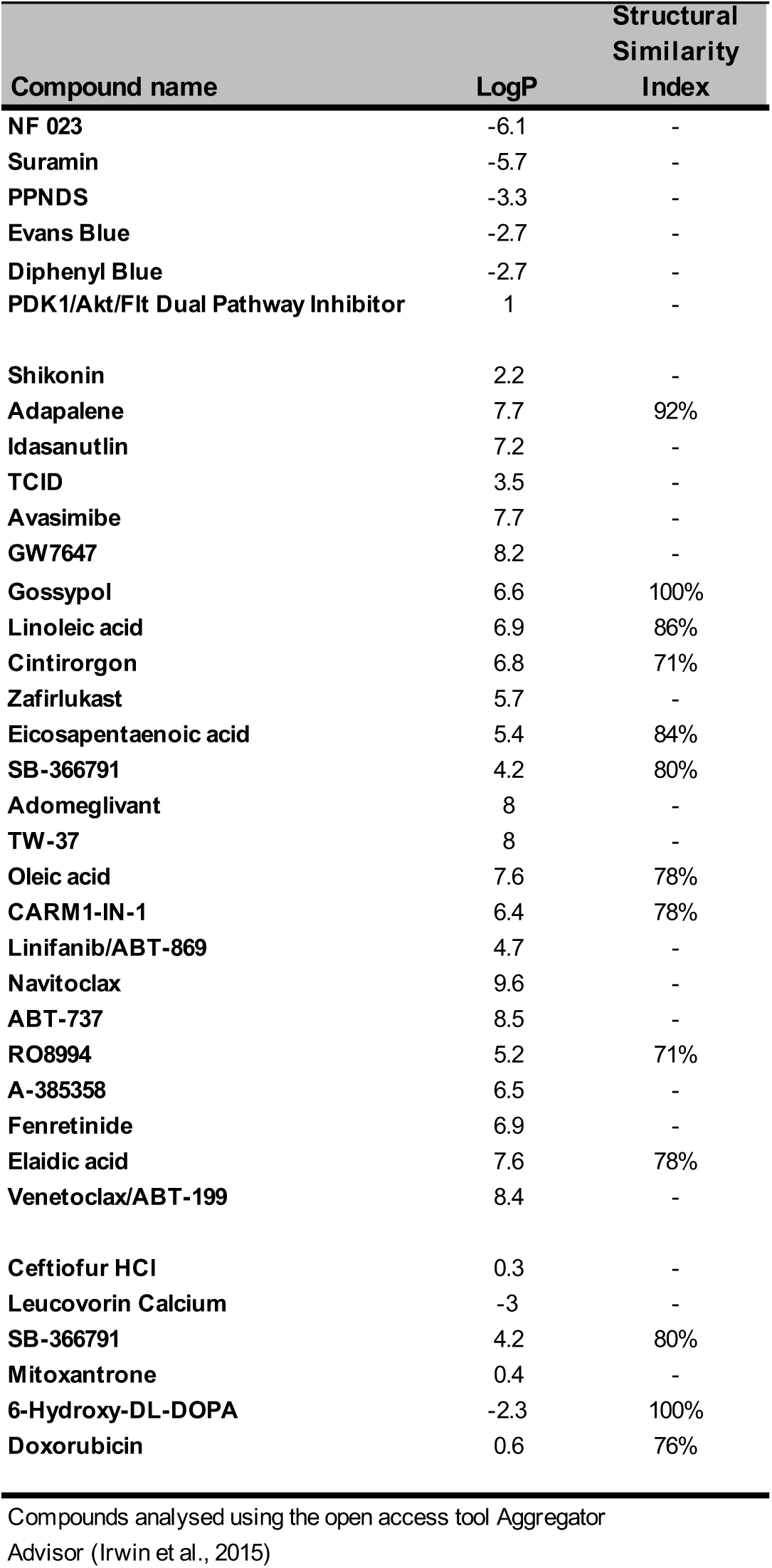
Predicted aggregation propensity of 35 HTS hit compounds selected in the first validation round

**Supplementary Table 3.** Predicted aggregation propensity of 12 HTS hit compounds selected in the second validation round and 2 control compounds

| Compound name | LogP | Structural Similarity Index |
| --- | --- | --- |
| Omapatrilat | 0.2 | - |
| Lifitegrast | 1.4 | - |
| FPA 124 | 1.7 | - |
| Eplerenone | 1.7 | 70% |
| Nedocromil | 2 | - |
| SKF 89145 hydrobromide | 2 | - |
| Fluocinolone Acetonide | 2.6 | - |
| NVP-BVU-972 | 3 | - |
| Hypocrellin A | 3.1 | - |
| ADL-5859 | 3.2 | - |
| TAK875 | 3.4 | - |
| Elvitegravir | 3.6 | - |
| **SSYA10-001<sup>a</sup>** | 2.1 | - |
| **Myricetin<sup>b</sup>** | 1.4 | 100% |

Compounds analysed using the open access tool Aggregator Advisor See (Irwin et al., 2015)
<sup>a</sup> Literature compound. See (Adedeji et al., Antimicrob Agents Chemother, 2012)
<sup>b</sup> Literature compound. See (Yu et al., Bioorg Med Chem Lett, 2012)

**Supplementary Table 4.**
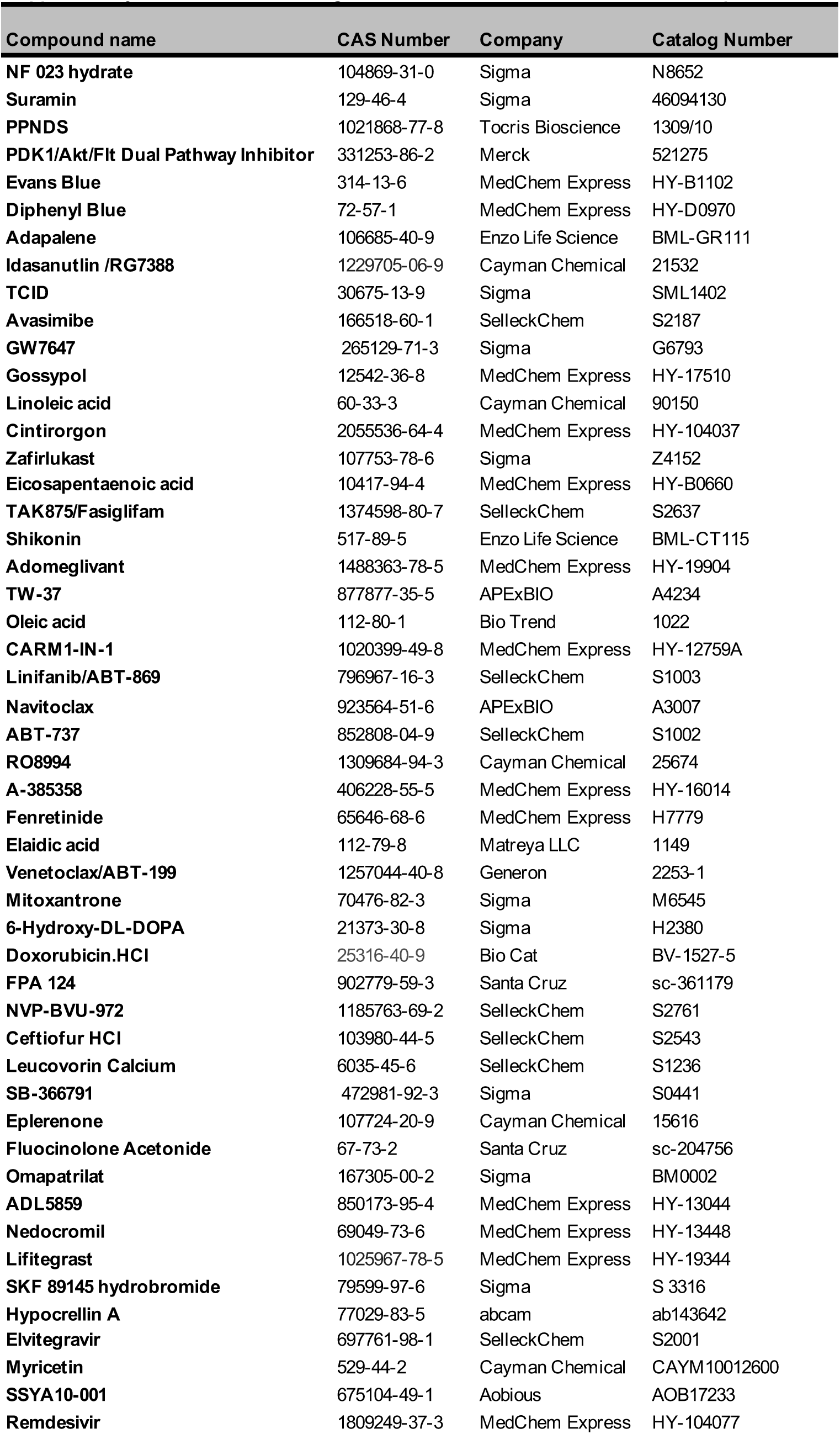
Purchased drugs for *in vitro* validation and viral inhibition experiments

